# Double reduction estimation and equilibrium tests in natural autopolyploid populations

**DOI:** 10.1101/2021.09.24.461731

**Authors:** David Gerard

## Abstract

Many bioinformatics pipelines include tests for equilibrium. Tests for diploids are well studied and widely available, but extending these approaches to autopolyploids is hampered by the presence of double reduction, the co-migration of sister chromatid segments into the same gamete during meiosis. Though a hindrance for equilibrium tests, double reduction rates are quantities of interest in their own right, as they provide insights about the meiotic behavior of autopolyploid organisms. Here, we develop procedures to (i) test for equilibrium while accounting for double reduction, and (ii) estimate double reduction given equilibrium. To do so, we take two approaches: a likelihood approach, and a novel *U*-statistic minimization approach that we show generalizes the classical equilibrium *χ*^2^ test in diploids. For small sample sizes and uncertain genotypes, we further develop a bootstrap procedure based on our *U*-statistic to test for equilibrium. Finally, we highlight the difficulty in distinguishing between random mating and equilibrium in tetraploids at biallelic loci. Our methods are implemented in the hwep R package on GitHub https://github.com/dcgerard/hwep.

## 1 Introduction

After its discovery, the Hardy-Weinberg law [Hardy, 1908, Weinberg, 1908] emerged as a foundational assumption in much of population genetics [Crow, 1988]. Since the implementation of the classical *χ*^2^ and likelihood ratio tests [Weir, 1996], the literature on testing the Hardy-Weinberg law in diploids has continued to grow [e.g. Levene, 1949, Haldane, 1954, Guo and Thompson, 1992, Wakefield, 2010, Graffelman and Weir, 2016, 2018, for a small sample] The research in polyploids is much more limited. In allopolyploids, if data from the different homoeologous genomes can be distinguished, then diploid approaches may be used. However, the research in equilibrium tests for autopolyploids is obstructed by the presence of double reduction, the co-migration of sister chromatid segments to the same gamete [Mather, 1935]. Double reduction increases homozygosity [Hardy, 2016] (Figure 1), introducing an inflation of Type I error into tests that ignore it (Section 3.1). It would therefore benefit the field to develop test procedures that account for double reduction.

**Figure 1:**
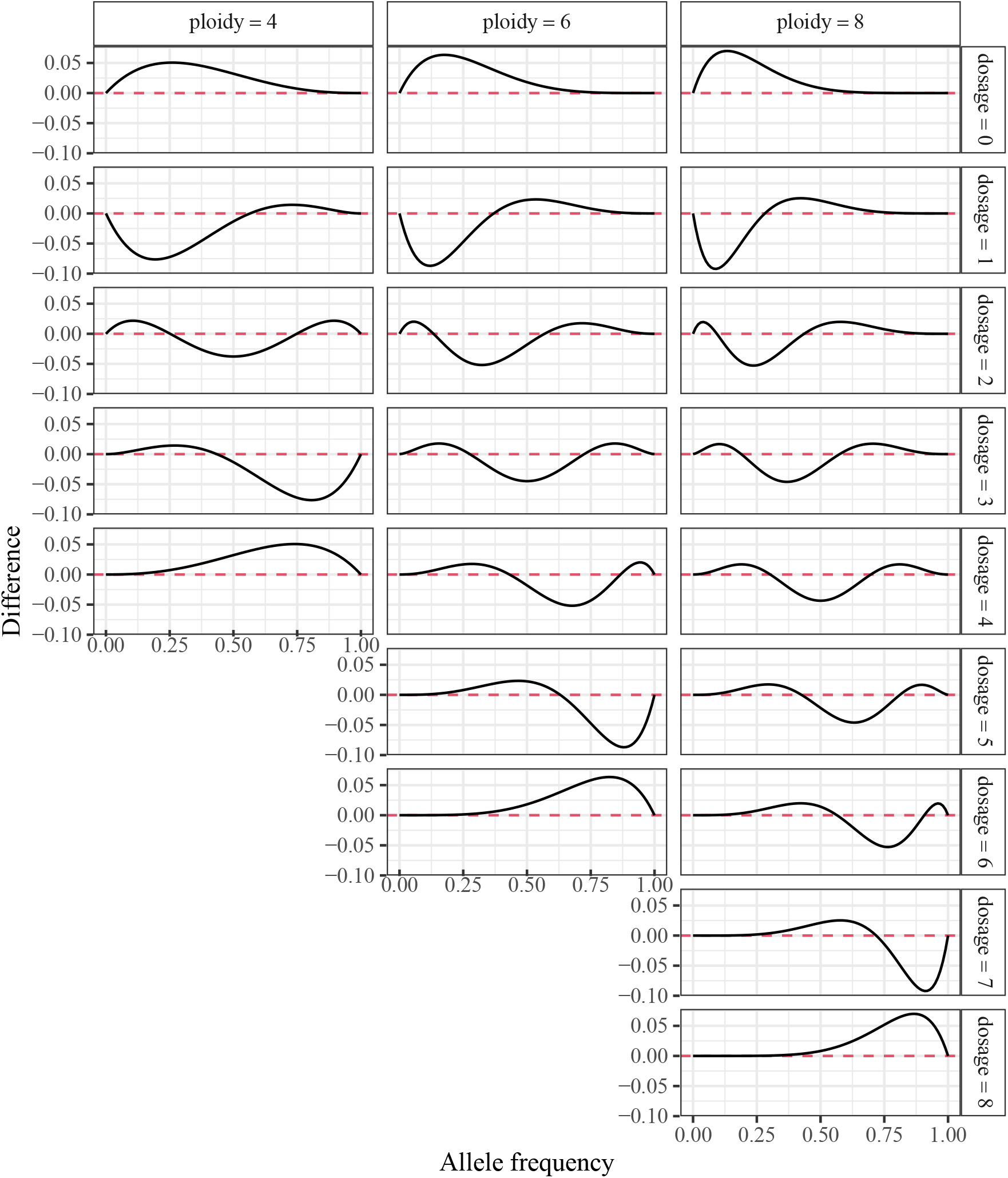
Over abundance of genotypes under the complete equational segregation model. The *y*-axis contains the equilibrium genotype frequencies under no double reduction subtracted from the equilibrium genotype frequencies under the complete equational segregation model [Mather, 1935]. The *x*-axis contains the allele frequency. Data are stratified by ploidy (column facets) and dosage (row facets). When the curves are above 0 (horizontal dashed line), this indicates that the dosage is overrepresented in the complete equational segregation model. Homozygosity is typically overrepresented, but the result for other dosages is more complex.

Estimating double reduction is also of interest in its own right due to its role in evolution [Meirmans et al., 2018], QTL mapping [Li et al., 2010], and plant breeding [Butruille and Boiteux, 2000], among other tasks. Most methods that estimate double reduction use experimental populations (such as F1 populations). Since experimental populations only require modeling a finite number of meioses for each offspring, this makes modeling much simpler. Using experimental populations, researchers have estimated double reduction either using segregation frequencies [Tai, 1982a,b, Haynes and Douches, 1993, Wu et al., 2001, Stift et al., 2008, Bourke et al., 2015], or they obtain estimates as a byproduct of haplotype reconstruction [Zheng et al., 2016, 2021]. For natural populations, the only manuscript we could find that claims to estimate double reduction is that of Jiang et al. [2021]. However, there is a vital flaw in the assumptions of this paper (Section S1), which results in biased estimates of double reduction (Section 3.1). In brief, Jiang et al. [2021] do not model equilibrium, but only one generation of possible double reduction.

If the equilibrium genotype frequencies, given double reduction, could be derived, then estimation and testing procedures would become feasible. Various manuscripts derive genotype frequencies at equilibrium in autopolyploids, but most without the full generality required to perform inference. Haldane [1930] provide genotype frequencies at equilibrium when there is no double reduction, but possible self-fertilization. Geiringer [1949] and Bennett [1968] provide genotype frequencies at equilibrium for tetraploids and hexaploids in terms of the rate of double reduction and the allele frequency of the population, but do not provide estimation or testing procedures, nor provide these frequencies at higher ploidy levels.

In a useful paper, Huang et al. [2019] layout a strategy for deriving genotype frequencies at equilibrium. One step in this strategy is to solve a complex system of nonlinear equations, a generically difficult task. Solving this system is possible for tetraploids (Section S3), and even for ploidies 6, 8, and 10 when using modern symbolic algebra systems (Section S4). But this approach becomes computationally infeasible for higher ploidies. Furthermore, the resulting equilibrium frequencies, because they are rather involved, provide no insight on any estimation and testing procedures that would use them, making this approach difficult to generalize to other models of meiosis or other evolutionary forces. And though Huang et al. [2019] derive these frequencies, they provide no means to test for equilibrium nor estimate double reduction given equilibrium.

In this manuscript, we will tackle equilibrium testing and double reduction estimation from multiple angles. In Section 2.3 we use the genotype frequencies derived by Huang et al. [2019] to implement a likelihood framework to test for equilibrium and estimate double reduction given equilibrium. However, because of the complexity of these genotype frequencies and the difficulty in obtaining such frequencies for larger ploidies, we develop a different approach for testing in Section 2.4. In this approach, we find a statistic that has expectation zero at equilibrium given the rate of double reduction. This motivates minimizing a norm of this statistic to make it as close to 0 as possible, thereby providing an estimate of the double reduction rate. We show that this minimized norm has an asymptotic *χ*^2^ distribution given equilibrium, which can be used to test for equilibrium. This approach generalizes the classical *χ*^2^ test for equilibrium in diploids to polyploids (Section S8). Finally, to account for small sample sizes and genotype uncertainty, in Section 2.5 we present a bootstrap approach to test for equilibrium.

There also exists a point of confusion in the literature on the difference between random mating and equilibrium. For example, Sun et al. [2020] develop a likelihood framework that they claim tests for equilibrium in tetraploids, but we would say actually tests for random mating (Section 2.2). In brief, we propose that random mating implies that genotype frequencies of the current generation are a convolution of the genotype frequencies of the gametes created by the parental generation, while equilibrium implies a steady state of genotype frequencies given a meiotic process and other population forces. We clarify this distinction in Section 2.1, and show that distinguishing between these hypotheses is rather subtle, particularly for tetraploids. The random mating hypothesis might be of interest, and so in Section 2.2 we generalize the approach of Sun et al. [2020] to higher ploidy levels.

## 2 Methods

### 2.1 Random mating versus equilibrium in tetraploids

In this section, we will highlight in tetraploids the difficulty of distinguishing between random mating and equilibrium while accounting for double reduction. In effect, our results show that it is impossible for any method to test for small deviations from equilibrium in the presence of double reduction using just a biallelic locus, but testing for moderate to large deviations from equilibrium is possible.

Genotype frequencies are structured under random mating. Let ***q*** = (*q*_0_,…, *q_K_*)^⊤^ denote the genotype frequencies of the *K* + 1 genotypes at a biallelic locus of a *K*-ploid species, where *K* is even. Consider the “gamete frequencies”, ***p*** = (*p*_0_,…, *p*_*K*/2_)^⊤^, where *p_i_* is the probability that a gamete has dosage *i*. Since individual genotypes are sums of gamete genotypes, random mating can be represented mathematically by

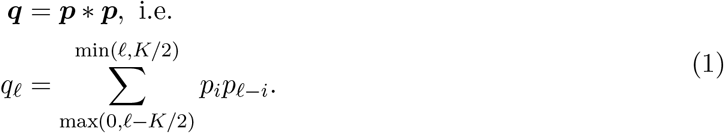

where “*” denotes discrete linear convolution. For example, in the case of diploids, we have ***p*** = (1 − *r, r*) for some allele frequency *r*, and ***q*** = ((1 − *r*)^2^, 2*r*(1 − *r*), *r*^2^)^⊤^, which is the well-known Hardy-Weinberg law.

On the other hand, equilibrium is a phenomenon that occurs after successive generations. To define equilibrium, we must first posit some process of updating genotype frequencies given the current genotype frequencies. Let *f* (***q, α***) be the updated genotype frequencies in the next generation given a rate of double reduction ***α*** = (*α*_1_,…, *α*_*K*/4_)^⊤^, where *α_i_* is the frequency that an individual contains *i* pairs of identical-by-double-reduction (IBDR) alleles (Section S5). Equilibrium occurs when the updated genotype frequencies are unchanged from one generation to the next, that is

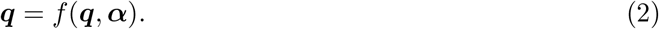

This update, *f*(·, ·), depends on the model for meiosis and other population genetic forces, random mating being one such possible force. The process we will use in this paper uses the same assumptions as used in deriving Hardy-Weinberg equilibrium in diploids [Waples, 2015]: (i) random mating, (ii) no selection, migration, gene flow, admixture, or mutation, (iii) only sexual selection, (iv) an infinite population size, (v) generations are non-overlapping, and (vi) allele frequencies are equal between sexes. The only generalization that we allow is double reduction.

As defined here, random mating is not the same as equilibrium. As noted by many authors, in polyploids it takes multiple generations of random mating to reach equilibrium [Haldane, 1930]. One clarifying example is that in an S1 population (one generation of selfing), the random mating hypothesis is satisfied, since the distribution of offspring genotypes is a convolution of the gamete probabilities within a single individual, but hardly any researcher would claim that the population is at equilibrium. However, Sun et al. [2020] conflated these hypotheses: they use null hypothesis (1) in tetraploids and claim that their resulting test is one against the null of equilibrium, when theirs in actuality is a test against the null of random mating.

Actually distinguishing between the random mating hypothesis and the equilibrium hypothesis is quite subtle. In tetraploids, there are two parameters under the random mating hypothesis (since *p*_0_ + *p*_1_ + *p*_2_ = 1), and there are two parameters under the equilibrium hypothesis (the double reduction rate, *α*, and the allele frequency, *r*). Thus, at equilibrium, there is a one-to-one correspondence between these two parameterizations (Section S3). In terms of ***p***, we have

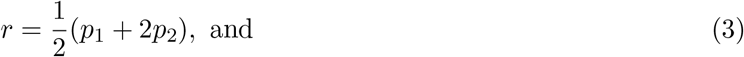

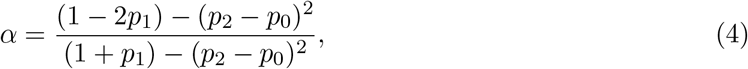

and in terms of (*α, r*), we have

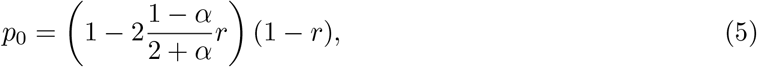

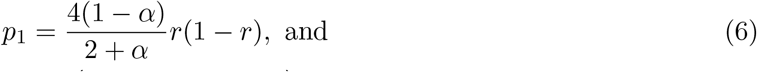

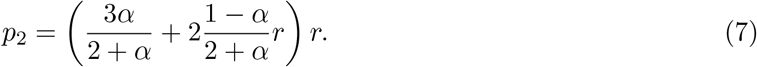

Because of this one-to-one correspondence, if we had random mating but *not* equilibrium, then the gamete frequencies ***p*** would correspond to *some* (*α, r*) at equilibrium, but we would not be able to tell if these are the actual values at equilibrium. Stated another way, because of this one-to-one correspondence, there are no left over degrees of freedom to distinguish between random mating and equilibrium.

Though there is a one-to-one correspondence between ***p*** and (*α, r*), not all values of ***p*** result in theoretically sound values of *α*. Other than constraining *α* to be non-negative, it is well known that under different theoretical models of meiosis the rate of double reduction is bounded above. E.g., under a complete equational segregation model, the maximum rate of double reduction is 1/6 [Mather, 1935]. If under equilibrium we have 0 ≤ *α* ≤ *c* for some *c* ∈ [0, 1], then this corresponds to the following gamete frequencies (derivation in Section S6).

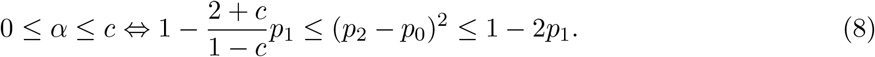

We can use this fact to test for extreme deviations from equilibrium, by testing if the gamete frequencies satisfy (8). However, even if the gamete frequencies satisfy (8), we are still unable to distinguish between random mating and equilibrium. It is only possible to test for moderate to major deviations from equilibrium.

A visualization of the region of gamete frequencies consistent with equilibrium is presented in Figure 2. If the gamete frequencies lie within this region, then they are consistent with *some* equilibrium, but we are not able to claim that they are *at* equilibrium. A summary of the nested hypotheses that can be tested at biallelic loci is presented in Table 1. We develop likelihood ratio tests to distinguish between these hypotheses in Section 2.3.

**Figure 2:**
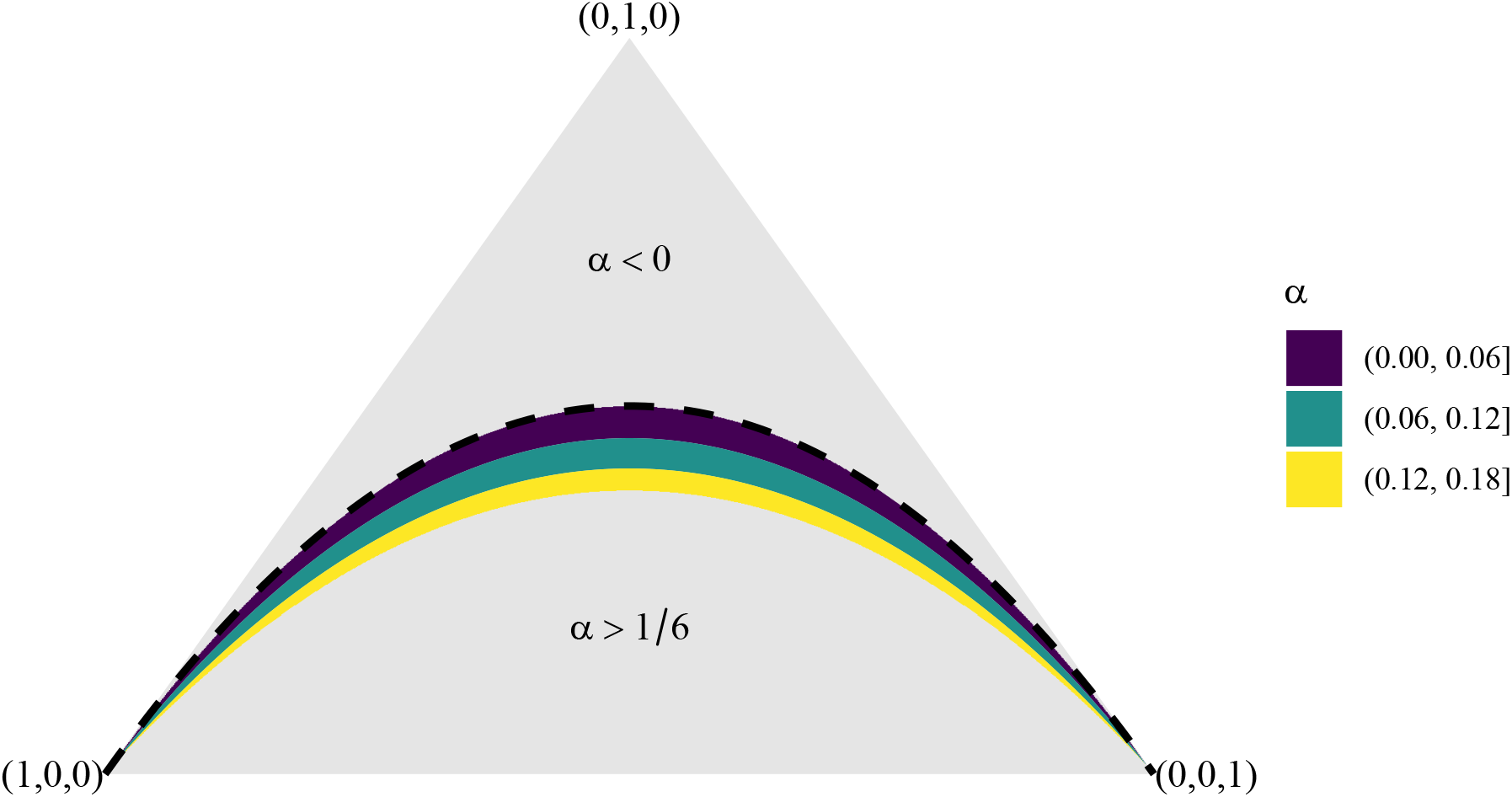
Isometric view of the unit 2-simplex, where each point on the triangle represents some combination of (*p*_0_, *p*_1_, *p*_2_). Colored bands are gametic frequencies consistent with some double reduction rate 0 ≤ *α* ≤ 1/6. Grey regions are inconsistent with double reduction rates within this interval. The dashed black line corresponds to *α* = 0. *H*_1_ (see Table 1 for a description of hypotheses) allows the gamete frequencies to lie anywhere in this simplex, *H*_2_ constrains the gamete frequencies to lie within the colored bands, and *H*_3_ constrains the gamete frequencies to lie on the dashed black line.

**Table 1:** Summary of hypotheses considered for tetraploids at biallelic loci.

| Hypothesis | Description | Parameter Constraints | Number of parameters |
| --- | --- | --- | --- |
| $H_0$ | Any genotype frequencies | $\sum_{i=0}^K q_i = 1$ | $K$ |
| $H_1$ | Random mating | $\mathbf{q} = \mathbf{p} * \mathbf{p}$ | $K/2$ |
| $H_2$ | Small deviations from equilibrium | $\mathbf{q} = \mathbf{p} * \mathbf{p}$ and<br>$1 - \frac{2+c}{1-c}p_1 \leq (p_2 - p_0)^2 \leq 1 - 2p_1$ | $\lfloor K/4 \rfloor + 1$ |
| $H_3$ | Equilibrium and no double reduction | $\mathbf{q} = \mathbf{p} * \mathbf{p}$ and<br>$\mathbf{p} = ((1-r)^2, 2r(1-r), r^2)$ | 1 |

An example of the difficulty in testing for equilibrium is graphically displayed in Figure 3, where we plot gamete frequencies along multiple generations of random mating. Generation 1 is an S1 population where the parental genotype was 2. The true level of *α* is 1/6. However, each iteration yields genotype frequencies that satisfy the random mating assumption (1), and thus correspond to *some* level of *α* that would maintain the gamete frequencies at their current levels. Only in generation 1 would this level of *α* be unreasonable, while the genotype frequencies are close to equilibrium only at around generation 5.

**Figure 3:**
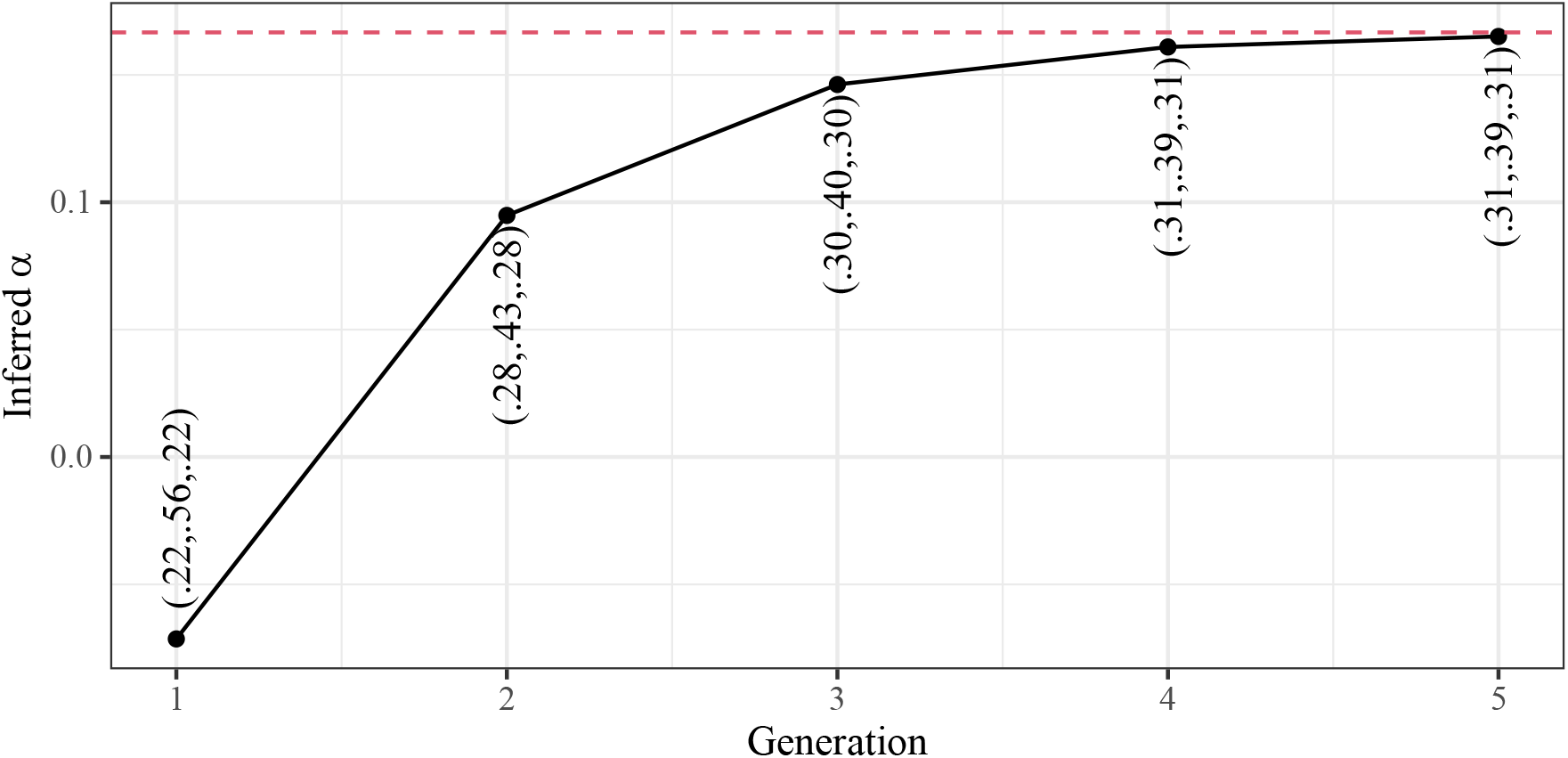
The inferred double reduction rate (*y*-axis) that would yield equilibrium for given gametes (tuples on plot) in successive generations (*x*-axis) of random mating when Generation 1 is an S1 population and the true double reduction rate is 1/6 (red dashed line).

### 2.2 Likelihood inference for random mating

In this section, we will generalize the likelihood framework of Sun et al. [2020] from tetraploids to general ploidy levels. Though Sun et al. [2020] claim that their likelihood approach is a test for equilibrium, as we have shown in Section 2.1, this is really a test for random mating. However, testing this hypothesis might be useful in its own right to clarify the mating behavior in a system.

To define a test against the null of random mating, let ***y*** = (*y*_0_,…, *y_K_*)^⊤^ denote the observed number of individuals with each genotype, so *y_i_* is the number of the individuals with genotype *i*. Let 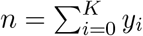 denote the total number of individuals. Then the likelihood is

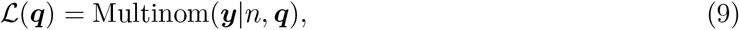

where Multinom(***y***|*n*, ***q***) denotes the probability mass function of the multinomial distribution with size *n* and probability vector ***q*** = (*q*_0_*,,…, q_K_*)^⊤^, evaluated at ***y***. Under the unconstrained model (where the population may or may not exhibit random mating), the maximum likelihood estimate of ***q*** is 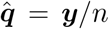. Under the constrained model where genotype frequencies are a result of random mating (1), we developed an expectation-maximization algorithm [Dempster et al., 1977] to maximize (9) using constraint (1) to obtain 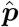 (Section S2). Under the null hypothesis of (1), the likelihood ratio test statistic,

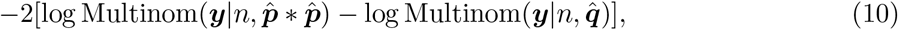

follows an asymptotic 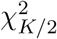 distribution, which we may use to calculate a *p*-value against the null of random mating.

### 2.3 Likelihood inference for equilibrium

In order to develop a likelihood framework to test against the null of equilibrium, we need to have access to the genotype frequencies at equilibrium given our meiotic process. Huang et al. [2019] developed a strategy to derive these genotype frequencies by setting up a system of non-linear equations. This system of equations is possible to solve in tetraploids (Section S3), and even in ploidies 6, 8, and 10 using modern symbolic algebra systems (Section S4). We will use these genotype frequencies to develop a likelihood ratio test for equilibrium. In tetraploids, this test is actually one against the null of at most minor deviations from equilibrium (Section 2.1).

Given a rate of double reduction ***α*** and an allele frequency *r*, we have access to the gamete frequencies at equilibrium ***p***. We will denote these gamete frequencies by ***p***(***α, r***) to emphasize their dependence on ***α*** and *r*. We can thus maximize the following likelihood (9) over 0 ≤ *α_i_* ≤ *c_i_* and *r* to obtain the MLEs,

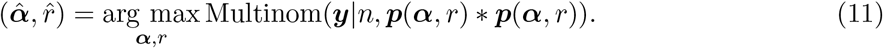

Then the likelihood ratio test statistic against the null of equilibrium is

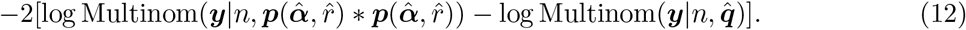

Equation (12) follows an asymptotic 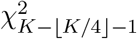 distribution under the null.

If one wanted to compare the null hypothesis of equilibrium against the alternative of random mating, one could calculate the following test statistic,

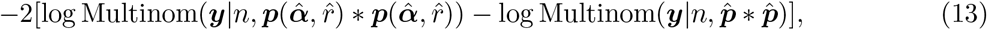

where 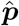 is calculated as in Section S2. The likelihood ratio test statistic (13) can be compared to a 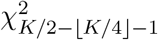 distribution, for ploidies greater than 4, to obtain a *p*-value. For tetraploids, we use a 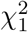 distribution, which is the asymptotic distribution of the test statistic under a fixed null value of *α* = 0 or *α* = *c*, a worst-case scenario for the test. Again, the null for tetraploids is one of only small deviations from equilibrium.

We can test against the null of no double reduction, given the population is in equilibrium. When the population is in equilibrium, but there is no double reduction, the genotypes follow a Binomial distribution with size *K* and success probability *r* [Haldane, 1930]. The MLE of *r* under the null is

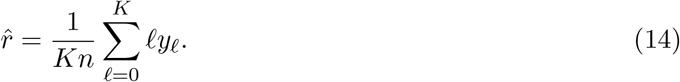

Let 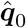 be the MLE of genotype frequencies at equilibrium given no double reduction, that is

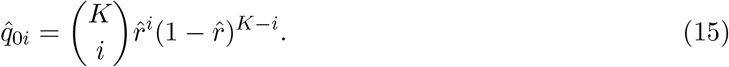

The test statistic is the following log-likelihood ratio

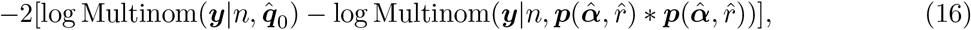

which under the null follows an asymptotic 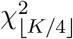 distribution, which may be used to calculate a *p*-value.

The asymptotics for all of these tests can be rather poor as many genotypes might exist with few or no individuals. To improve the asymptotic approximation (at the risk of losing power), we apply an aggregation strategy. Specifically, we aggregate genotypes that have no individuals, adjusting the resulting degrees of freedom of the *χ*^2^ distributions accordingly.

Though we were able to use the genotype frequencies at equilibrium, these formulas are rather complicated, sometimes spanning pages. They thus provide no insight behind the methods used to test for equilibrium. This makes it difficult to generalize the likelihood approach to other models of meiosis or population forces. Furthermore, for higher ploidies, the system of non-linear equations becomes computationally infeasible, at least given the author’s computational resources, and so these methods are difficult to generalize to higher ploidy individuals. All of this motivates the development of an alternative procedure that is more intuitive. We will present such an approach in Section 2.4.

### 2.4 A *U*-statistic approach to estimate double reduction and test for equilibrium given a meiotic process

In this Section we will develop a novel approach to test for equilibrium and to estimate double reduction given equilibrium, implemented for every ploidy level. We derive a statistic, called a “*U*-statistic” [Hoeffding, 1948], that has expectation zero at equilibrium given the rate of double reduction. To obtain an estimate of this rate, we minimize a norm of the *U*-statistic over the double reduction rate. We can show that, at equilibrium, this minimized norm has an asymptotic *χ*^2^ distribution, which we can use to obtain a *p*-value for a test against the null of equilibrium. This method is a generalization to polyploids, in the presence of double reduction, of the classical *χ*^2^ test for equilibrium in diploids (Section S8). Formal details of our method are presented in Section S7. Here, we will provide a more intuitive description.

Let us begin by deriving a statistic that has expectation zero at equilibrium. Let *Pr*(*g*|*G*_1_, *G*_2_, ***θ***) be the probability of two parents with genotypes *G*_1_, *G*_2_ ∈ {0, 1,…, *K*} producing an offspring with genotype *g* ∈ {0, 1,…, *K*}. This probability depends on the model of meiosis, and we denote the parameters of this meiotic model by ***θ***. For this manuscript, we use the model for meiosis in Section S5, and so ***θ*** = ***α***, the double reduction rate. But our methods in this section would applicable to different models of meiosis. Let *x_i_*, *x_j_*, and *x_ℓ_* be the genotypes for individuals *i*, *j*, and *ℓ*. Consider the following statistic

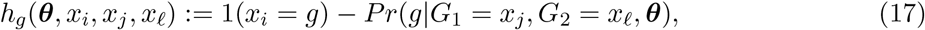

where 1(·) is the indicator function. The idea behind (17) is that we are choosing two individuals (*j* and *ℓ*) to act as the “parents” to the “child” (*i*). We then compare the probability of child *i* having genotype *g* to whether or not child *i* actually has genotype *g*.

Since at equilibrium individuals *j* and *ℓ* will be identically distributed to the actual parents of individual *i*, it is easy to show that (17) has expectation zero at equilibrium,

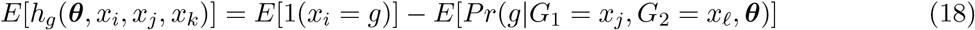

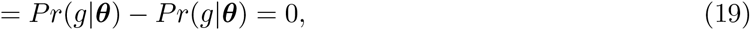

where *Pr*(*g*|***θ***) is the frequency of genotype *g* at equilibrium. The *h_g_*(***θ***, *x_i_, x_j_, x_k_*) statistic (17) only involves three individuals, but we can average over all possible triples of individuals to come up with a statistic that still has (approximately) expectation zero.

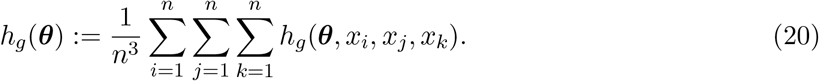

Equation (20) is our “*U*-statistic” [Hoeffding, 1948]. (Technically, (20) is actually a “*V*-statistic” [von Mises, 1947], but these have the same asymptotic behavior as *U*-statistics, and since *U*-statistics are more familiar to the Statistics community, we will stick to this terminology). Equation (20) actually represents *K* + 1 *U*-statistics for *g* = 0, 1,…, *K*, each having expectation zero at equilibrium. We’ll let the vector of these *U*-statistics be represented by

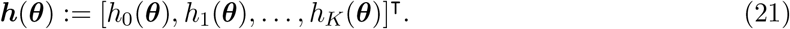

Due to sampling variability, the *h_g_*(***θ***)’s will not actually equal zero. But because they should equal zero in expectation, one intuitive way to estimate ***θ*** is to find a ***θ*** that makes (21) as close to **0** as possible. We do this by minimizing a norm of ***h***(***θ***),

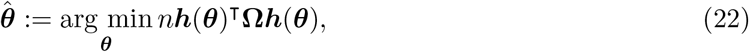

where **Ω** is some positive semi-definite matrix. Such estimators are sometimes called “*U*-process minimizers” or “*M_m_* estimators” in the literature [Honoré and Powell, 1994, Bose, 2002, Bose and Chatterjee, 2018]. It turns out that, for a properly chosen value of **Ω**, the norm evaluated at 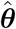,

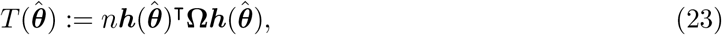

has an asymptotic *χ*^2^ distribution under the null of equilibrium. This result is derived from the theory of generalized method of moments from the econometrics literature [Hansen, 1982, Newey and West, 1987], and is an application of the Sargan-Hansen *J* test. We can use this result to explicitly test for equilibrium.

There is another intuitive way to describe (21). Let ***q*** = (*q*_0_, *q*_1_,…, *q_K_*)^⊤^ represent the genotype frequencies of a population. Then, under random mating, and given a model for meiosis, there is a function *f* (·, ·) that updates the genotype frequencies from one generation to the next. That is, the genotype frequencies of the following generation would be *f* (***q, θ***). The specific form of *f* (·, ·) used in our manuscript is described in Section S5, but our methods are more general. If we let 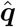 be the empirical genotype frequencies of a sample, then one can show that

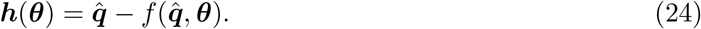

That is, our vector of *U*-statistics is the difference between the empirical genotype frequencies and one update of the empirical genotype frequencies to the next generation. This is very intuitive, as at equilibrium we have ***q*** = *f* (***q, θ***) (2), and so we would expect 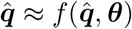. This formulation also shows that we can calculate the *U*-statistic in linear time, instead of cubic time, since the most computationally intensive part is calculating the sample frequencies 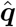. However, framing (24) in terms of *U*-statistics allows us to wield the asymptotic machinery developed by the field of Statistics, and derive a null distribution given equilibrium.

The full estimation algorithm is presented in Procedure 1. See Section S7 for a formal derivation of our results, Section S7.1 for some practical methodological considerations, and Section S8 for a proof that our *U*-statistic approach generalizes the classical *χ*^2^ test from diploids to polyploids.

### 2.5 A bootstrap approach for small samples and uncertain genotypes

Two deficiencies arise from from the methods of Sections 2.3 and 2.4. First, these approaches require known genotypes. Genotype uncertainty is a severe issue in polyploid genetics research [Gerard et al., 2018, Gerard and Ferrão, 2019], motivating approaches to account for it, such as when estimating linkage disequilibrium [Gerard, 2021a,b] or inbreeding [Blischak et al., 2018]. The ubiquity of genotype uncertainty limits the applicability of the methods in Sections 2.3 and 2.4 (though, these methods appear to be conservative when using posterior mean genotype counts, see Section S11). Second, these approaches are heavily dependent on asymptotic approximations, so might not be accurate for small sample sizes. Exact tests (or Monte Carlo approximations to exact tests) are generally preferable to those requiring asymptotic approximations [Rohlfs and Weir, 2008]. In polyploids, permutation approaches have been created in the SPAGeDi [Hardy and Vekemans, 2002] and GENODIVE [Meirmans and Van Tienderen, 2004] softwares. However, because the null distribution these programs use is that under random segregation of alleles, this approach effectively tests for the presence of binomial genotype frequencies, being unable to account for double reduction.

To overcome both of these deficiencies, we propose a bootstrap approach [Efron, 1979] applied to our *U*-statistic. Let *c_ik_* denote the posterior probability that individual *i* = 1, 2,…, *n* has genotype *k* = 0, 1,…, *K*. These posterior probabilities are obtainable from many genotyping programs [Voorrips et al., 2011, Serang et al., 2012, Gerard et al., 2018, Gerard and Ferrão, 2019]. If there were no genotype uncertainty, then *c_ik_* would be an indicator function for the true genotype for individual *i*. One bootstrap iteration consists of first sampling *n* individuals *with replacement*, then sampling each sampled individual’s genotype using the *c_ik_*’s, then estimating ***α*** by minimizing norm (22), then finally calculating a test statistic. If 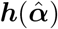 is the vector of observed *U*-statistics (21), and 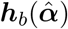 is the *b*th bootstrap replication of this vector of *U*-statistics, then we propose to use the following test statistic

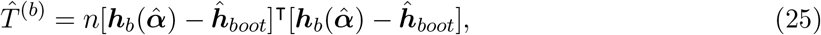

where 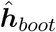 is the empirical mean of the bootstrap 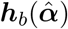’s. We repeat this process for *B* bootstrap replications. To obtain a *p*-value against the null of equilibrium, we calculate the proportion of 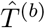’s that are larger than

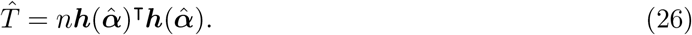

The intuition is that the bootstrap samples of (25) approximate the distribution of the distance of 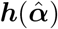 from its mean, so we can compare the distance of 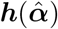 from **0**, calculated in (26), to this bootstrap distribution to see if **0** is a reasonable value for the mean. Details of the algorithm are presented in Procedure 2.

## 3 Results

### 3.1 Simulations

We ran simulations to study the accuracy of the double reduction estimators, and to evaluate the properties of the proposed tests for equilibrium. Each replication, we generated genotypes of *n* ∈ {25, 100, 1000} individuals of ploidy *K* ∈ {4, 6, 8}, given the major allele frequency *r* ∈ {0.1, 0.25, 0.5} and the double reduction parameters ***α*** ∈ {**0**, ***α**_m_*/2, ***α**_m_*}, where ***α***_*m*_ contains the maximum values of the double reduction parameters under the complete equational segregation model (S76). Given *r* and ***α*** we generated allele frequencies ***q*** after several rounds of random mating using Procedure 3. The number of rounds of random mating was used as a proxy for deviation from equilibrium, where more rounds of random mating result in frequencies closer to equilibrium. If *m* is the maximum number of rounds of random mating in Procedure 3, we set *m* ∈ {1, 2, 3, ∞}, where *m* = 1 results in the furthest deviation from equilibrium and *m* = ∞ results in the frequencies at equilibrium. Each replication, given the generated ***q***, we simulated the number of individuals with each genotype ***y*** ~ Multinom(*n, **q***). We ran 1000 replications for each unique combination of parameter settings.

Given these simulated counts, estimates for ***α*** were obtained using the likelihood approach (Section 2.3), the *U*-statistic approach (Section 2.4), and (for tetraploids) the method of Jiang et al. [2021]. These simulated counts were also used to run the *U*-statistic test for equilibrium (Section 2.4), the likelihood ratio test for equilibrium (Section 2.3), the likelihood ratio test for random mating (Section 2.2), the bootstrap test for equilibrium (Section 2.5), as well as a likelihood ratio test for equilibrium that does not account for double reduction (Section S10). Due to computational demands, the bootstrap method was only run for a subset of the simulations for only 200 replications each, specifically for *n* = 25, *K* = 8, *r* ∈ {0.1, 0.5}, *α* ∈ {**0**, ***α**_m_*/2, ***α**_m_*}, and *m* ∈ {1, 3, ∞}.

The results for estimating ***α*** are presented in Figures S2–S4. There, we see that the method of Jiang et al. [2021] is highly biased. Our methods are somewhat biased for small sample sizes, but appear to be consistent for larger sample sizes. The likelihood and the *U*-statistic approaches appear to behave very similarly, with the likelihood approach being slightly biased up and the *U*-statistic approach being slightly biased down for small sample sizes. The variability of the estimators is quite large, even for larger sample sizes, highlighting the difficulty in estimating the rate of double reduction using just a single biallelic locus.

Test statistics of the *U*-statistic approach and the likelihood approach both generally follow their theoretical distributions under the null, except when the allele frequency is small for tetraploids (Figures S5–S10). The bootstrap approach also produces uniform *p*-values when the null is satisfied (Figure S11). All three tests are mostly able to control type I error at the 0.05 level, being only slightly anti-conservative in some settings, while the classical approach (Section S10) is completely unable to control type I error except when there is no double reduction (Figure S12–S14). The *U* - statistic, likelihood, and bootstrap approaches behave very similarly in terms of power, appearing to be asymptotically equivalent (Figures S15–S17). All three can distinguish between the null and alternative hypotheses, but have small power for only small deviations from equilibrium. To understand the power results for tetraploids in S15–S17, recall from Section 2.1 that all sets of gamete frequencies have an “inferred” double reduction rate at which these gamete frequencies would indeed be at equilibrium. We plot this inferred rate in Figure S18 and we see that after three generations, the *α* = 0 and *α* = *α_m_*/2 scenarios have inferred rates within the bounds of theoretically sound double reduction rates. Thus, after three generations of random mating, these are also null scenarios, as they only exhibit small deviations from equilibrium.

All scenarios we simulated under satisfy the random mating assumption, and so we also explored the ability of the random mating test of Section 2.2 to control Type I error in Figures S19–S21. The test is generally able to control for Type I error, being only somewhat anti-conservative for small sample sizes.

See Section S11 for an initial exploration on the effects of gentoype uncertainty.

### 3.2 Analysis of a hatchery population of *Acipenser transmontanus*

In this section, we analyze the dataset of white sturgeon (*Acipenser transmontanus*) from Delomas et al. [2021]. These data consist of ancestral octoploids that likely exhibit tetrasomic inheritance, with disomic inheritance being likely rare in sturgeon [Drauch Schreier et al., 2011]. These data were collected from a Central California caviar farm with broodstock from the Sacramento River, and so likely exhibit random mating and possibly only minor deviations from equilibrium caused, for example, by unobserved relatedness.

We took the *n* = 19 tetraploid individuals from these data and applied updog [Gerard et al., 2018, Gerard and Ferrão, 2019] on the sequencing data to obtain genotype estimates. These data were sequenced to an exceptionally high depth, with a mean average read-depth of 5342, and so it is reasonable to take these genotypes as known. We then tested for equilibrium using the likelihood approach (Section 2.3), the *U*-statistic approach (Section 2.4), the bootstrap approach (Section 2.5), and the approach that assumes no double reduction (Section S10). We also tested for random mating (Section 2.2).

The random mating test generally produced more uniform *p*-values than the other tests, while the equilibrium approaches that allow for double reduction generally produced more uniform-looking *p*-values than the likelihood ratio test that does not account for double reduction (Figures S22–S23). One can partially explain this by the excess homozygosity we observe in some loci. For example, Figure S24 explores two loci where the *p*-values assuming no double reduction were 0.058 and 0.056 while the *p*-values using the *U*-statistic approach were 0.32 and 0.33. There, we plot the observed genotype counts against the expected genotype counts while either allowing for double reduction, or assuming no double reduction. We see that there is an excess of homozygosity, which can be partially accounted for by modeling double reduction.

We do not display the resulting estimates of double reduction as simulations indicate that such estimates are highly unreliable for such a small sample size (Section 3.1).

### 3.3 Analysis of an S1 population of *Ipomoea batatas*

We evaluated our methods on an S1 population autohexaploid sweetpotato (*Ipomoea batatas*, 2*n* = 6*x* = 90), obtained from Shirasawa et al. [2017]. SNPs were filtered using the `VariantAnnotation` package [Obenchain et al., 2014] using the following criteria: allele frequencies between 0.1 and 0.9, average read depth at least 100, and no more than 50% missing data. We obtained genotype estimates for each individual at each locus with `updog` [Gerard et al., 2018, Gerard and Ferrão, 2019]. Genotype counts were then fed into `hwep` to test for equilibrium using the likelihood approach (Section 2.3), the *U*-statistic approach (Section 2.4), and the naive approach (Section S10). We did not run the bootstrap approach due to the computational burden of this method. These data are not at equilibrium, and so each method is expected to produce small *p*-values. We also tested for random mating (Section 2.2), which should be fulfilled in these data.

Our results are presented in Figures S25–S27. We see that all equilibrium tests behave similarly and result in small *p*-values (Figure S25). The test for random mating is very conservative (Figure S26), likely because each locus has a large number of genotypes with no or few individuals, worsening the asymptotic approximation of this approach. However, the random mating method provides accurate estimates of gamete frequencies, which we compared to the theoretical hypergeometric distribution of gamete frequencies given the estimated parental genotype at each locus (Figure S27).

## 4 Discussion

In this manuscript, we clarified the distinction between random mating and equilibrium, noting subtleties in distinguishing between these two hypotheses in the presence of double reduction. We then developed likelihood approaches to both test for random mating and test for equilibrium. Our likelihood approach for equilibrium also yields an estimate of double reduction given equilibrium. However, because of the complexity of genotype frequencies at equilibrium, we developed an intuitively simpler approach based on a statistic that is, on average, zero at equilibrium. This simpler approach reduces down to the classical *χ*^2^ test for Hardy-Weinberg equilibrium in diploids. For small samples and uncertain genotypes, we developed a bootstrap approach to test for equilibrium that uses our *U*-statistic formulation. We verified our methods in simulations and on real data.

Double reduction has a similar effect on equilibrium as inbreeding [Hardy, 2016]. Thus, it is possible that any double reduction estimates might be picking up the effects of inbreeding. However, one benefit of our approach is that if one rejects the null of equilibrium, then this might suggest that greater inbreeding has occurred than can be accounted for by double reduction, while an approach that does not account for double reduction would not be able to make this claim. It might be possible to adjust our *U*-statistic approach to incorporate inbreeding. For example, instead of choosing individuals uniformly to be the “parents” of “child” *i* in (17), one could incorporate a parameter that makes individual *i* more likely to be its own “parent”. Having an additional parameter would make the model unidentified, at least in tetraploids. However, because inbreeding is likely a individual-specific phenomenon [Blischak et al., 2018] while double reduction is a locus-specific phenomenon, this might inspire an estimation strategy for these parameters. We do not pursue these ideas further in this manuscript.

Our methods do not allow for mixed ploidy populations. Such populations are of interest, as they allow for the study of the evolutionary consequences of different ploidy levels [McAllister and Miller, 2016, Monnahan et al., 2019, Wos et al., 2019], or they exist as a consequence of fish management [Delomas, 2019]. One benefit of the *U*-statistic approach is that it would be feasible to modify it for mixed ploidy populations with special forms of meiosis, such as those found in *Andropogon gerardii* [Norrmann et al., 1997]. For example, if individual *i* had ploidy *K_i_*, and we had a model for meiosis given parental genotypes and ploidies, say *Pr*(*g*|*G*_1_ = *x_j_*, *G*_2_ = *x_ℓ_, K_j_, K_ℓ_, **θ***), then we could develop a *U*-statistic for evaluating equilibrium in this mixed-ploidy population, based on the following,

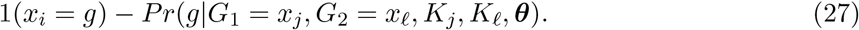

We leave studying and implementing this extension for future work.

Our methods also do not allow for odd ploidy levels. Though rare, all-triploid bisexual reproduction is possible [Stöck et al., 2002]. The equilibrium frequencies for such species have not yet been derived, limiting the ability to apply a likelihood approach to test for equilibrium. However, our *U*-statistic approach would potentially provide a strategy to test for equilibrium, as long as a model for meiosis was provided.

## Acknowledgments

This material is based upon work supported by the National Science Foundation under Grant No. 2132247.

We express our thanks to Dr. Dennis Leung, University of Melbourne, and Dr. Y. Samuel Wang, Cornell University, for insightful discussions. Many thanks to Dr. Thomas Delomas, Pacific States Marine Fisheries Commission, for help with the *Acipenser transmontanus* data.

Most analyses were performed using the R statistical language [R Core Team, 2020].

## Data availability

The hwep R package implementing the methods described in this paper is available on GitHub (https://github.com/dcgerard/hwep). All scripts required to reproduce the results of this manuscript are available at https://github.com/dcgerard/hwesims. All datasets used in this manuscript are open and available at https://www.doi.org/10.5061/dryad.crjdfn33r and http://sweetpotato-garden.kazusa.or.jp/.

## 5 Tables, figures, and procedures

**Procedure 1.**
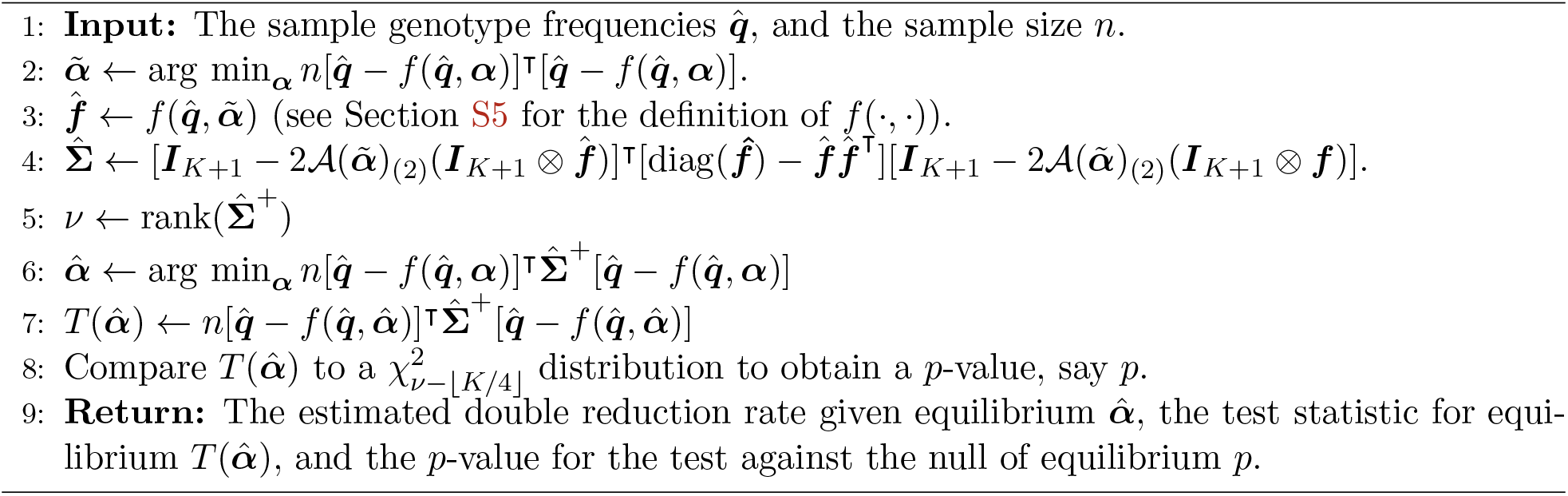
Procedure to estimate ***α*** given equilibrium, and test for equilibrium while accounting for double reduction. 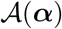 is a three-dimensional array such that 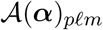 is the probability an offspring will have genotype *p* given parental genotypes *ℓ* and *m* (Section S5), 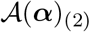 is the second-mode matricization of 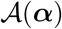 [Kolda and Bader, 2009], ⊗ is the Kronecker product, and 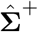 is the Moore-Penrose inverse of 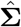. See Section S7 for a derivation.

**Procedure 2.**
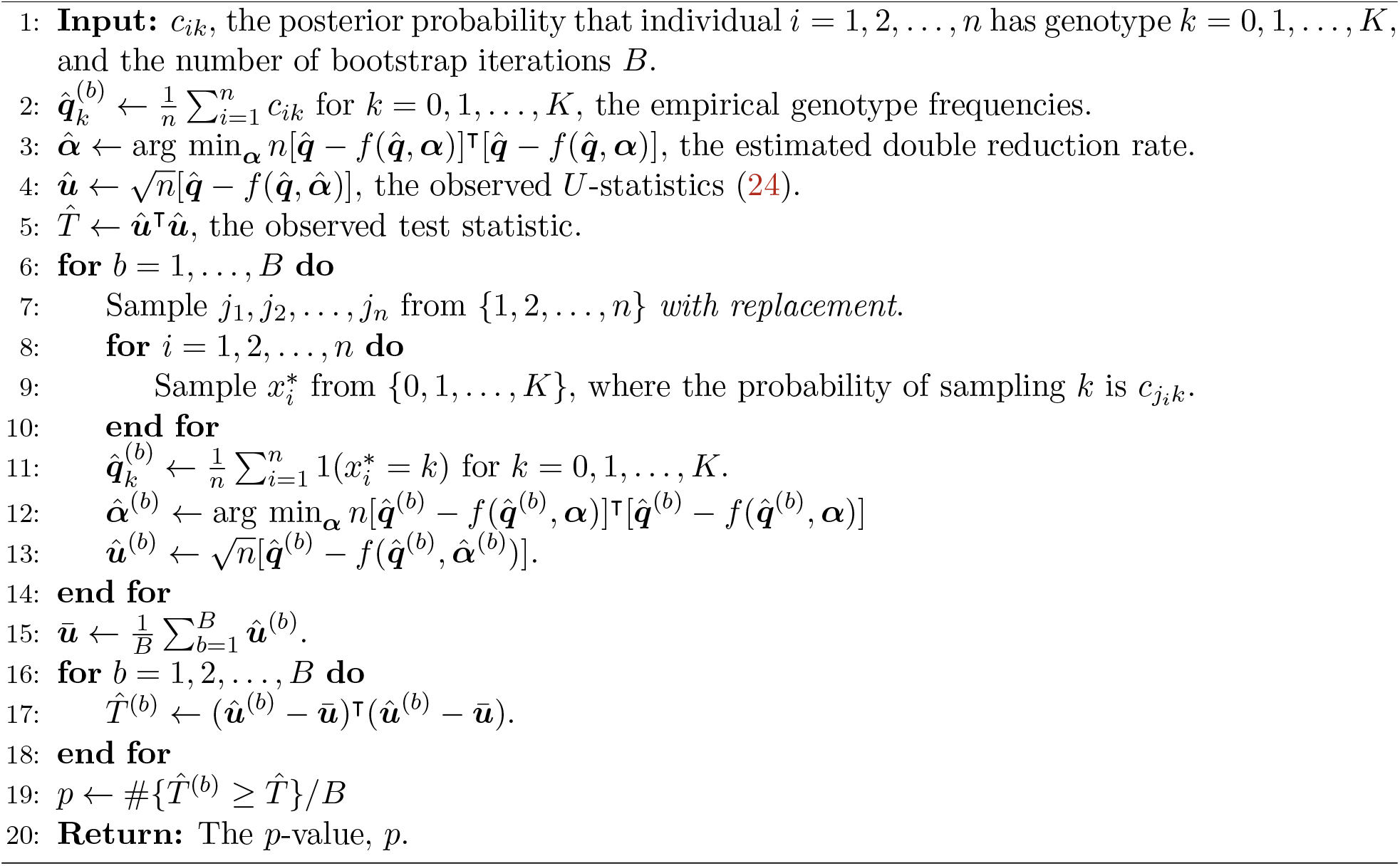
Bootstrap procedure for calculating a *p*-value against the null of equilibrium while accounting for both genotype uncertainty and small sample sizes.

**Procedure 3.**
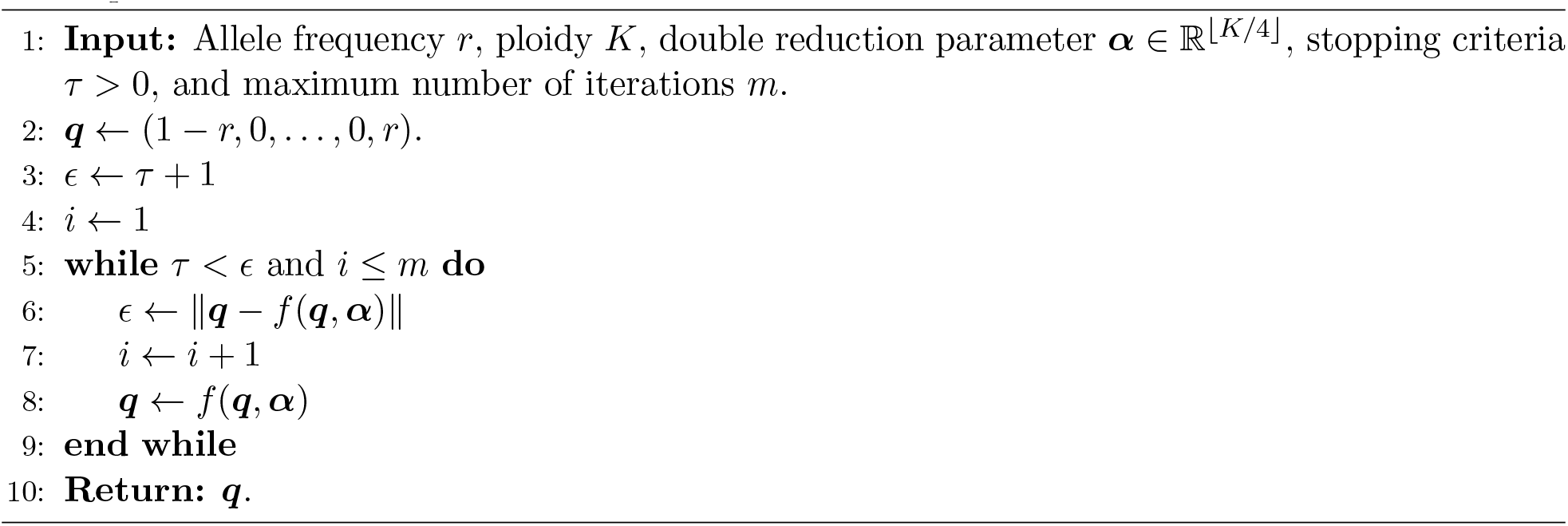
Procedure to generate allele frequencies after several rounds of random mating given the allele frequency *r*, double reduction parameter ***α***, and ploidy *K*. Allele frequencies are typically near equilibrium after five iterations.

