## Supplementary Material for "Double reduction estimation and equilibrium tests in natural autopolyploid populations"

David Gerard

Department of Mathematics and Statistics, American University, Washington, DC, 20016, USA

**Abstract**

This file contains supplementary figures, details, and theoretical considerations for "Double reduction estimation and equilibrium tests in natural autopolyploid populations".

#### S1 Flaw of Jiang et al. [2021]

The vital flaw occurs with Equation (5) from Jiang et al. [2021]. There, they assume that the gametic frequencies of the grandparent generation are, using our notation,

$$\begin{aligned} p_2 &= r^2 \\ p_1 &= 2r(1-r) \\ p_0 &= (1-r)^2, \end{aligned} \tag{S1}$$

for some allele frequency $r \in [0, 1]$. However, since the population is at equilibrium this would mean that the parent genotype frequencies are

$$\begin{aligned} q_4 &= p_2^2 = r^4 \\ q_3 &= 2p_2p_1 = 4r^3(1-r) \\ q_2 &= 2p_2p_0 + p_1^2 = 6r^2(1-r)^2 \\ q_1 &= 2p_1p_0 = 4r(1-r)^3 \\ q_0 &= p_0^2 = (1-r)^4, \end{aligned} \tag{S2}$$

Thus, an implicit assumption of Jiang et al. [2021] is that the parent dosages are Binomially distributed with size 4 and success probability of $r$. This is the distribution of genotypes at equilibrium under no double reduction. They then update these genotype frequencies using one iteration of what we do in Section S5, with double reduction, to come up with a model of the genotype frequencies in the current generation. As noted by Sun et al. [2020], it takes more than 6 generations for a tetraploid species to be close to equilibrium, and so the genotype frequencies of Jiang et al. [2021] are mispecified (Figure S1).

#### S2 EM algorithm to estimate gamete frequencies under random mating

In this section, we derive an EM algorithm to maximize likelihood (9) under constraint (1).

Let $x_i \in \{0, \ldots, K\}$ denote the genotype for individual $i$. Let $Z_{ijk}$ be the latent variable that denotes the parental genotypes for individual $i$, where

$$Z_{ijk} = \begin{cases} 1 & \text{if } j \leq k \text{ and one parent has genotype } j \text{ and the other has genotype } k \\ 0 & \text{otherwise} \end{cases} \tag{S3}$$

Then the complete log-likelihood is

$$\sum_{i=1}^{n} \sum_{j=\max(0, x_i - K/2)}^{\lfloor x_i/2 \rfloor} Z_{i,j,x_i-j}[\log(p_j) + \log(p_{x_i-j}) + 1(j \neq x_i/2)\log(2)], \tag{S4}$$

where $1(\cdot)$ is the indicator function. The E-step corresponds to calculating the following quantities

$$w_{j\ell} := E[Z_{i,j,x_i-j}|x_i = \ell] = \frac{p_j p_{\ell-j} 2^{1(j\neq \ell/2)}}{\sum_{j=\max(0,\ell-K/2)}^{\lfloor \ell/2 \rfloor} p_j p_{\ell-j} 2^{1(j\neq\ell/2)}}. \tag{S5}$$

The M-step involves maximizing the following over $\boldsymbol{p}$:

$$\sum_{i=1}^{n} \sum_{j=\max(0, x_i - K/2)}^{\lfloor x_i/2 \rfloor} E[Z_{i,j,x_i-j}|x_i][\log(p_j) + \log(p_{x_i-j}) + 1(j \neq x_i/2)\log(2)] \tag{S6}$$

$$= \sum_{\ell=0}^{K} \sum_{j=\max(0,\ell-K/2)}^{\lfloor \ell/2 \rfloor} y_\ell w_{j\ell}[\log(p_j) + \log(p_{\ell-j})] + \text{constant}, \tag{S7}$$

where $y_\ell = \sum_{i=1}^{n} 1(x_i = \ell)$ is the number of individuals with genotype $\ell$. Collecting the coefficients in (S7) together, we get that (S7) is equal to

$$\sum_{m=0}^{K/2} \eta_m \log(p_m) + \text{constant}, \text{ where} \tag{S8}$$

$$\eta_m := \sum_{j=0}^{m} w_{jm} y_{j+m} + \sum_{k=m}^{K/2} w_{mk} y_{m+k}. \tag{S9}$$

Equation (S8) is proportional to a multinomial log-likelihood. Thus, the update for $\boldsymbol{p}$ is

$$p_m = \frac{\eta_m}{\sum_{j=0}^{K/2} \eta_j}. \tag{S10}$$

The EM algorithm consists of iteratively calculating the $w_{j\ell}$'s in (S5) and updating the $p_m$'s with (S9) and (S10).

#### S3 Correspondence between gamete frequencies and double reduction in tetraploids at equilibrium

Here, we will derive the one-to-one correspondence between the gamete frequencies $(p_0, p_1, p_2)$ and $(\alpha, r)$, where $\alpha$ is the probability of double reduction and $r$ is the allele frequency of one allele.

Under the random mating hypothesis, we have

$$
\begin{aligned}
q_0 &= p_0^2, && \text{(S11)}\\
q_1 &= 2p_0p_1, && \text{(S12)}\\
q_2 &= 2p_0p_2 + p_1^2, && \text{(S13)}\\
q_3 &= 2p_1p_2, \text{ and} && \text{(S14)}\\
q_4 &= p_2^2. && \text{(S15)}
\end{aligned}
$$

Assuming the population is at equilibrium, using the the segregation probabilities from Fisher and Mather [1943], and using the law of total probability, we have that

$$
\hat{p}_0 = \hat{q}_0 + (\frac{1}{2} + \frac{1}{4}\hat{\alpha})\hat{q}_1 + (\frac{1}{6} + \frac{1}{3}\hat{\alpha})\hat{q}_2 + \frac{1}{4}\hat{\alpha}\hat{q}_3. \quad \text{(S16)}
$$

Using equations (S11)–(S15) we can solve for $\hat{\alpha}$ in terms of the gamete frequencies. After simplifying, using the fact that $\hat{p}_0 + \hat{p}_1 + \hat{p}_2 = 1$, we obtain (4).

To solve for $(p_0, p_1, p_2)$ in terms of $(\alpha, r)$, we use (4) along with the following two equations

$$
\begin{aligned}
r &= \frac{1}{2}(p_1 + 2p_2) && \text{(S17)}\\
1 &= p_0 + p_1 + p_2, && \text{(S18)}
\end{aligned}
$$

to set up a system of three equations and three unknowns. Solving, we get (5)–(7).

See Bennett [1968] for a different technique of obtaining these frequencies.

#### S4 Gamete frequencies at equilibrium for higher ploidies

We used Mathematica [Wolfram Research, Inc., 2020] to obtain genotype frequencies at equilibrium for hexaploids (see online material `https://github.com/dcgerard/hwesims`). To do so, we solve the following system of equations for $\boldsymbol{p}$, and simplify the resulting forms

$$
\begin{aligned}
\boldsymbol{p} &= \boldsymbol{B}(\boldsymbol{\alpha})^\intercal(\boldsymbol{p} * \boldsymbol{p}), && \text{(S19)}\\
r &= \frac{1}{K/2}\sum_{i=0}^{K/2} ip_i, && \text{(S20)}
\end{aligned}
$$

where $\boldsymbol{B}(\boldsymbol{\alpha})$ is defined in Section S5. We obtained

$$
p_0 = \left(1 - \frac{9(3-\alpha)(6-\alpha)}{(9+\alpha)(9+2\alpha)}r + \frac{27(1-\alpha)(3-\alpha)}{(9+\alpha)(9+2\alpha)}r^2\right)(1-r) \quad \text{(S21)}
$$

$$p_1 = \left( \frac{9(3-\alpha)(9-4\alpha)}{(9+\alpha)(9+2\alpha)} - \frac{81(1-\alpha)(3-\alpha)}{(9+\alpha)(9+2\alpha)} r \right) r(1-r) \tag{S22}$$

$$p_2 = \left( \frac{45\alpha(3-\alpha)}{(9+\alpha)(9+2\alpha)} + \frac{81(1-\alpha)(3-\alpha)}{(9+\alpha)(9+2\alpha)} r \right) r(1-r) \tag{S23}$$

$$p_3 = \left( \frac{20\alpha^2}{(9+\alpha)(9+2\alpha)} + \frac{45\alpha(3-\alpha)}{(9+\alpha)(9+2\alpha)} r + \frac{27(1-\alpha)(3-\alpha)}{(9+\alpha)(9+2\alpha)} r^2 \right) r \tag{S24}$$

For ploidies 8 and 10, Huang et al. [2019] derived the genotype frequencies in terms of the double reduction rates and the allele frequency. They did so also using a symbolic algebra system with the system of equations (S19)–(S20). The resulting equations are quite involved, and so we do not present them here, as they provide no intuitive understanding.

#### S5 Updating genotype frequencies

We will describe how to update genotypic frequencies to the next generation given current genotypic frequencies. We do so using the F1 segregation probabilities developed by Fisher and Mather [1943] for tetraploids and hexaploids, and then later generalized in Huang et al. [2019] to all ploidy levels. To describe these segregation probabilities in terms of a biallelic loci, let the random variable $G \in \{0, \ldots, K\}$ be the dosage of an individual with even ploidy $K$, with distribution $\boldsymbol{q} := (q_0, \ldots, q_K)$ where $\sum_{i=0}^{K} q_i = 1$. Let the random variable $g \in \{0, \ldots, K/2\}$ be the dosage of a gamete produced by that individual. Let $\alpha_i$ be the probability that a gamete carries $i$ copies of identical-by-double-reduction (IBDR) alleles, for $i = 0, \ldots, \lfloor K/4 \rfloor$ and $\sum_{i=0}^{\lfloor K/4 \rfloor} \alpha_i = 1$, where $\lfloor \cdot \rfloor$ is the floor function. Then, for biallelic loci, Equation (1) from Huang et al. [2019] reduces to:

$$Pr(g|G, \boldsymbol{\alpha}) = \sum_{i=0}^{\lfloor K/4 \rfloor} \sum_{j=0}^{i} \frac{\binom{G}{j}\binom{G-j}{g-2j}\binom{K-G}{i-j}\binom{K-G-(i-j)}{K/2-g-2(i-j)}}{\binom{K}{i}\binom{K-i}{K/2-2i}} \alpha_i. \tag{S25}$$

Let $\boldsymbol{B}(\boldsymbol{\alpha}) \in \mathbb{R}^{(K+1)\times(K/2+1)}$ contain the probabilities in (S25), where $\boldsymbol{B}(\boldsymbol{\alpha})_{ij} = Pr(g = j | G = i, \boldsymbol{\alpha})$. We call this the segregation matrix, since it contains the segregation probabilities of gamete dosages given parental dosages. Then, by the law of total probability, the marginal probability of gamete dosages can be represented by

$$\boldsymbol{p}(\boldsymbol{q}, \boldsymbol{B}(\boldsymbol{\alpha})) = \boldsymbol{B}(\boldsymbol{\alpha})^{\intercal} \boldsymbol{q}, \tag{S26}$$

where $\boldsymbol{p}(\boldsymbol{B}(\boldsymbol{\alpha}))_i$ is the marginal probability of a gamete having dosage $i$. The updated genotype frequencies are a discrete linear convolution of $\boldsymbol{p}(\boldsymbol{B}(\boldsymbol{\alpha}))$ with itself:

$$f(\boldsymbol{q}, \boldsymbol{\alpha}) = \boldsymbol{p}(\boldsymbol{q}, \boldsymbol{B}(\boldsymbol{\alpha})) * \boldsymbol{p}(\boldsymbol{q}, \boldsymbol{B}(\boldsymbol{\alpha})). \tag{S27}$$

Equation (S27) represents updating genotype frequencies in terms of gamete segregation probabilities. An alternative, but equivalent, way of representing genotype frequency updates is in terms of *zygote* segregation probabilities. Let $\boldsymbol{B}(\boldsymbol{\alpha})_{[j,]}$ represent the $j$th row of $\boldsymbol{B}(\boldsymbol{\alpha})$, and let $\mathcal{A}(\boldsymbol{\alpha}) \in \mathbb{R}^{(K+1)\times(K+1)\times(K+1)}$ be a multidimensional array such that

$$\mathcal{A}(\boldsymbol{\alpha})_{ijk} = (\boldsymbol{B}(\boldsymbol{\alpha})_{[j,]} * \boldsymbol{B}(\boldsymbol{\alpha})_{[k,]})_i. \tag{S28}$$

That is, $\mathcal{A}(\boldsymbol{\alpha})_{ijk}$ is the probability an offspring will have genotype $i$ when the parents have genotypes $j$ and $k$. Then, using the law of total probability, we have

$$f(\boldsymbol{q}, \boldsymbol{\alpha})_i = \sum_{j=0}^{K} \sum_{k=0}^{K} \mathcal{A}(\boldsymbol{\alpha})_{ijk} q_j q_k. \tag{S29}$$

We can represent this in terms of the Tucker product [Kolda and Bader, 2009] by

$$f(\boldsymbol{q}, \boldsymbol{\alpha}) = (\boldsymbol{I}_{K+1}, \boldsymbol{q}^\intercal, \boldsymbol{q}^\intercal) \cdot \mathcal{A}(\boldsymbol{\alpha}). \tag{S30}$$

#### S6 Derivation of equation (8)

Using (4), we have that the denominator is always greater than 0 (since $1 + p_1 \geq 1 \geq (p_2 - p_0)^2$), and denominator is always greater than the numerator. Thus, $\alpha \geq 0$ if an only if the numerator is negative, if and only if $(p_2 - p_0)^2 \leq 1 - 2p_1$.

Again using (4), we have

$$\alpha < c \tag{S31}$$

$$\frac{(1 - 2p_1) - (p_2 - p_0)^2}{(1 + p_1) - (p_2 - p_0)^2} \leq c \tag{S32}$$

$$\Leftrightarrow (1 - 2p_1) - (p_2 - p_0)^2 \leq c + cp_1 - c(p_2 - p_0)^2 \tag{S33}$$

$$\Leftrightarrow (1 - c) - (2 + c)p_1 \leq (1 - c)(p_2 - p_0)^2 \tag{S34}$$

$$\Leftrightarrow 1 - \frac{2 + c}{1 - c} p_1 \leq (p_2 - p_0)^2. \tag{S35}$$

#### S7 Estimating double reduction and testing for equilibrium

In this section, we will derive a method to test for equilibrium in the presence of double reduction, while also estimating the degree of double reduction given equilibrium. Our strategy is to derive a $U$-statistic that has expectation zero if the population is at equilibrium. This will motivate a procedure to minimize a norm of this $U$-statistic to estimate the degree of double reduction. At equilibrium, this minimized norm follows a $\chi^2$ distribution, which will allow us to test for equilibrium while accounting for double reduction.

Let $\boldsymbol{e}_j$ denote the $j$th unit vector in $K + 1$ dimensions, where the $j$th element of $\boldsymbol{e}_j$ is 1, and every other element is 0. Let $\boldsymbol{Y}_i$ be a random variable which equals $\boldsymbol{e}_j$ if individual $i$ has genotype $j$. The $\boldsymbol{Y}_i$'s are assumed to be independent with $Pr(\boldsymbol{Y}_i = \boldsymbol{e}_j) = q_j$. Let $\mathcal{A}(\boldsymbol{\alpha}) \in \mathbb{R}^{(K+1)\times(K+1)\times(K+1)}$ be a multidimensional array such that $\mathcal{A}(\boldsymbol{\alpha})_{p\ell m}$ is the probability an offspring will have genotype $p$ given parental genotypes $\ell$ and $m$ (Section S5).

**Lemma S1.** *Let $1 \leq i, j, k \leq n$ with $j \neq k$. Then, at equilibrium,*

$$E[\boldsymbol{Y}_i - (\boldsymbol{I}_{K+1}, \boldsymbol{Y}_j^\intercal, \boldsymbol{Y}_k^\intercal) \cdot \mathcal{A}(\boldsymbol{\alpha})] = \boldsymbol{0}. \tag{S36}$$

*Proof.* We have $E[\boldsymbol{Y}_i] = \boldsymbol{q}$. We also have, by linearity of the Tucker product and the independence between $\boldsymbol{Y}_j$ and $\boldsymbol{Y}_k$,

$$
\begin{aligned}
E[(\boldsymbol{I}_{K+1}, \boldsymbol{Y}_j^\intercal, \boldsymbol{Y}_k^\intercal) \cdot \mathcal{A}(\boldsymbol{\alpha})] &= (\boldsymbol{I}_{K+1}, E[\boldsymbol{Y}_j]^\intercal, E[\boldsymbol{Y}_k]^\intercal) \cdot \mathcal{A}(\boldsymbol{\alpha}) && \text{(S37)} \\
&= (\boldsymbol{I}_{K+1}, \boldsymbol{q}^\intercal, \boldsymbol{q}^\intercal) \cdot \mathcal{A}(\boldsymbol{\alpha}) && \text{(S38)} \\
&= f(\boldsymbol{q}, \boldsymbol{\alpha}). && \text{(S39)}
\end{aligned}
$$

Where equality (S39) results from (S30). Since at equilibrium we have $\boldsymbol{q} = f(\boldsymbol{q}, \boldsymbol{\alpha})$, the result is proved. □

Thus, given $\boldsymbol{\alpha}$ is known, the following is a $U$-statistic [Van der Vaart, 2000] that has expectation $\mathbf{0}$ at equilibrium:

$$
\hat{U}_n(\boldsymbol{\alpha}) := \frac{1}{\binom{n}{3}} \sum_{1 \leq i < j < k \leq n} \boldsymbol{h}(\boldsymbol{Y}_i, \boldsymbol{Y}_j, \boldsymbol{Y}_k | \boldsymbol{\alpha}), \text{ where,} \tag{S40}
$$

$$
\begin{aligned}
\boldsymbol{h}(\boldsymbol{Y}_i, \boldsymbol{Y}_j, \boldsymbol{Y}_k | \boldsymbol{\alpha}) =& [\boldsymbol{Y}_i - (\boldsymbol{I}_{K+1}, \boldsymbol{Y}_j^\intercal, \boldsymbol{Y}_k^\intercal) \cdot \mathcal{A}(\boldsymbol{\alpha}) + \\
& \boldsymbol{Y}_j - (\boldsymbol{I}_{K+1}, \boldsymbol{Y}_i^\intercal, \boldsymbol{Y}_k^\intercal) \cdot \mathcal{A}(\boldsymbol{\alpha}) + \\
& \boldsymbol{Y}_k - (\boldsymbol{I}_{K+1}, \boldsymbol{Y}_i^\intercal, \boldsymbol{Y}_j^\intercal) \cdot \mathcal{A}(\boldsymbol{\alpha})]/3.
\end{aligned} \tag{S41}
$$

However, instead of working with $U$-statistic (S40), we will instead work with the closely related, but conceptually much simpler, $V$-statistic

$$
\hat{V}_n(\boldsymbol{\alpha}) := \frac{1}{n^3} \sum_{i,j,k} V(\boldsymbol{Y}_i, \boldsymbol{Y}_j, \boldsymbol{Y}_k | \boldsymbol{\alpha}) \text{ where} \tag{S42}
$$

$$
V(\boldsymbol{Y}_i, \boldsymbol{Y}_j, \boldsymbol{Y}_k | \boldsymbol{\alpha}) := \boldsymbol{Y}_i - (\boldsymbol{I}_{K+1}, \boldsymbol{Y}_j^\intercal, \boldsymbol{Y}_k^\intercal) \cdot \mathcal{A}(\boldsymbol{\alpha}). \tag{S43}
$$

which has the same asymptotic behavior as its $U$-statistic counterpart [Zhou et al., 2019]. It turns out that this $V$-statistic is equivalent to the difference between the current genotype frequencies, and the genotype frequencies after one update of Section S5 (Theorem S1).

**Theorem S1.**

$$
\begin{aligned}
\hat{V}_n(\boldsymbol{\alpha}) &= \frac{1}{n^3} \sum_{i,j,k} [\boldsymbol{Y}_i - (\boldsymbol{I}_{K+1}, \boldsymbol{Y}_j^\intercal, \boldsymbol{Y}_k^\intercal) \cdot \mathcal{A}(\boldsymbol{\alpha})] && \text{(S44)} \\
&= \frac{1}{n} \sum_{i=1}^{n} (\boldsymbol{Y}_i - f(\hat{\boldsymbol{q}}, \boldsymbol{\alpha})) && \text{(S45)} \\
&= \hat{\boldsymbol{q}} - f(\hat{\boldsymbol{q}}, \boldsymbol{\alpha}), && \text{(S46)}
\end{aligned}
$$

*where* $\hat{\boldsymbol{q}} = \frac{1}{n} \sum_{i=1}^{n} \boldsymbol{Y}_i$.

*Proof.* It is clear that

$$
\frac{1}{n^3} \sum_{i,j,k} \boldsymbol{Y}_i = \frac{1}{n} \sum_{i=1}^{n} \boldsymbol{Y}_i = \hat{\boldsymbol{q}}. \tag{S47}
$$

By the definition of the Tucker product [Kolda and Bader, 2009], we have

$$\begin{aligned}
\frac{1}{n^3}\sum_{ijk}[(\boldsymbol{I}_{K+1}, \boldsymbol{Y}_j^\intercal, \boldsymbol{Y}_k^\intercal)\cdot \mathcal{A}(\boldsymbol{\alpha})]_p &= \frac{1}{n^3}\sum_{ijk}\sum_{\ell=0}^{K}\sum_{m=0}^{K}\mathcal{A}(\boldsymbol{\alpha})_{p\ell m}\boldsymbol{Y}_{j\ell}\boldsymbol{Y}_{km} && \text{(S48)}\\
&= \frac{1}{n^2}\sum_{j=1}^{n}\sum_{k=1}^{n}\sum_{\ell=0}^{K}\sum_{m=0}^{K}\mathcal{A}(\boldsymbol{\alpha})_{p\ell m}\boldsymbol{Y}_{j\ell}\boldsymbol{Y}_{km} && \text{(S49)}\\
&= \sum_{\ell=0}^{K}\sum_{m=0}^{K}\mathcal{A}(\boldsymbol{\alpha})_{p\ell m}\left(\frac{1}{n}\sum_{j=1}^{n}\boldsymbol{Y}_{j\ell}\right)\left(\frac{1}{n}\sum_{k=1}^{n}\boldsymbol{Y}_{km}\right) && \text{(S50)}\\
&= \sum_{\ell=0}^{K}\sum_{m=0}^{K}\mathcal{A}(\boldsymbol{\alpha})_{p\ell m}\hat{\boldsymbol{q}}_\ell\hat{\boldsymbol{q}}_m && \text{(S51)}\\
&= [(\boldsymbol{I}_{K+1}, \hat{\boldsymbol{q}}^\intercal, \hat{\boldsymbol{q}}^\intercal)\cdot \mathcal{A}(\boldsymbol{\alpha})]_p. && \text{(S52)}
\end{aligned}$$

Thus $\frac{1}{n^3}\sum_{ijk}(\boldsymbol{I}_{K+1}, \boldsymbol{Y}_j^\intercal, \boldsymbol{Y}_k^\intercal)\cdot \mathcal{A}(\boldsymbol{\alpha}) = (\boldsymbol{I}_{K+1}, \hat{\boldsymbol{q}}^\intercal, \hat{\boldsymbol{q}}^\intercal)\cdot \mathcal{A}(\boldsymbol{\alpha}) = f(\hat{\boldsymbol{q}}, \boldsymbol{\alpha})$, by (S30). □

For our estimation and testing procedures, we will need the asymptotic covariance matrix of $\sqrt{n}(\hat{V}_n(\boldsymbol{\alpha}))$, provided by Theorem S2.

**Theorem S2.** *At equilibrium, we have*

$$\operatorname{cov}[\sqrt{n}\hat{V}_n(\boldsymbol{\alpha})] \to [\boldsymbol{I}_{K+1} - 2\mathcal{A}(\boldsymbol{\alpha})_{(2)}(\boldsymbol{I}_{K+1}\otimes \boldsymbol{q})]^\intercal \boldsymbol{Q}[\boldsymbol{I}_{K+1} - 2\mathcal{A}(\boldsymbol{\alpha})_{(2)}(\boldsymbol{I}_{K+1}\otimes \boldsymbol{q})], \tag{S53}$$

*where $\otimes$ is the Kronecker product, $\mathcal{A}(\boldsymbol{\alpha})_{(2)}$ is the second-mode matricization of $\mathcal{A}(\boldsymbol{\alpha})$ [Kolda and Bader, 2009], and*

$$\boldsymbol{Q} := \operatorname{diag}(\boldsymbol{q}) - \boldsymbol{q}\boldsymbol{q}^\intercal = \begin{pmatrix} q_0(1-q_0) & -q_0q_1 & \cdots & -q_0q_K \\ -q_0q_1 & q_1(1-q_1) & \cdots & -q_1q_K \\ \vdots & \vdots & \ddots & \vdots \\ -q_0q_K & -q_1q_K & \cdots & q_K(1-q_K)\end{pmatrix}. \tag{S54}$$

*Proof.* We will actually derive the asymptotic covariance of (S40), which has the same asymptotic covariance as (S42). Standard methods for $U$ statistics state that [Van der Vaart, 2000]

$$\operatorname{cov}(\hat{U}_n(\boldsymbol{\alpha})) \approx \frac{9}{n}\operatorname{cov}[\boldsymbol{h}(\boldsymbol{X}, \boldsymbol{Y}_2, \boldsymbol{Y}_3), \boldsymbol{h}(\boldsymbol{X}, \boldsymbol{Z}_2, \boldsymbol{Z}_3)] \tag{S55}$$

where $\boldsymbol{X}, \boldsymbol{Y}_2, \boldsymbol{Y}_3, \boldsymbol{Z}_2, \boldsymbol{Z}_3$ are *iid*. Plugging in $\boldsymbol{h}(\cdot,\cdot,\cdot)$ from (S41) into (S55), we get (note that the 9 in (S55) cancels with the two 1/3's in (S41))

$$\begin{aligned}
&\operatorname{cov}(\sqrt{n}\hat{U}_n(\boldsymbol{\alpha}))\\
&\approx \operatorname{cov}[\boldsymbol{X} + \boldsymbol{Y}_1 + \boldsymbol{Y}_2 -\\
&\qquad (\boldsymbol{I}_{K+1}, \boldsymbol{Y}_1^\intercal, \boldsymbol{Y}_2^\intercal)\cdot\mathcal{A}(\boldsymbol{\alpha}) - (\boldsymbol{I}_{K+1}, \boldsymbol{X}^\intercal, \boldsymbol{Y}_2^\intercal)\cdot\mathcal{A}(\boldsymbol{\alpha}) - (\boldsymbol{I}_{K+1}, \boldsymbol{X}^\intercal, \boldsymbol{Y}_3^\intercal)\cdot\mathcal{A}(\boldsymbol{\alpha}),\\
&\qquad \boldsymbol{X} + \boldsymbol{Z}_1 + \boldsymbol{Z}_2 -\\
&\qquad (\boldsymbol{I}_{K+1}, \boldsymbol{Z}_1^\intercal, \boldsymbol{Z}_2^\intercal)\cdot\mathcal{A}(\boldsymbol{\alpha}) - (\boldsymbol{I}_{K+1}, \boldsymbol{X}^\intercal, \boldsymbol{Z}_2^\intercal)\cdot\mathcal{A}(\boldsymbol{\alpha}) - (\boldsymbol{I}_{K+1}, \boldsymbol{X}^\intercal, \boldsymbol{Z}_3^\intercal)\cdot\mathcal{A}(\boldsymbol{\alpha})]
\end{aligned} \tag{S56}$$

$$
\begin{aligned}
&= \operatorname{cov}[\boldsymbol{X}, \boldsymbol{X}] - 2\operatorname{cov}[\boldsymbol{X}, (\boldsymbol{I}_{K+1}, \boldsymbol{X}^\intercal, \boldsymbol{Y}^\intercal) \cdot \mathcal{A}(\boldsymbol{\alpha})] - 2\operatorname{cov}[(\boldsymbol{I}_{K+1}, \boldsymbol{X}^\intercal, \boldsymbol{Y}^\intercal) \cdot \mathcal{A}(\boldsymbol{\alpha}), \boldsymbol{X},] \\
&\quad + 4\operatorname{cov}[(\boldsymbol{I}_{K+1}, \boldsymbol{X}^\intercal, \boldsymbol{Y}^\intercal) \cdot \mathcal{A}(\boldsymbol{\alpha}), (\boldsymbol{I}_{K+1}, \boldsymbol{X}^\intercal, \boldsymbol{Z}^\intercal) \cdot \mathcal{A}(\boldsymbol{\alpha})] \quad \text{(S57)} \\
&= \operatorname{cov}[\boldsymbol{X}, \boldsymbol{X}] - 2\operatorname{cov}[\boldsymbol{X}, (\boldsymbol{I}_{K+1} \otimes \boldsymbol{Y}^\intercal)\mathcal{A}(\boldsymbol{\alpha})_{(2)}^\intercal \boldsymbol{X}] - 2\operatorname{cov}[(\boldsymbol{I}_{K+1} \otimes \boldsymbol{Y}^\intercal)\mathcal{A}(\boldsymbol{\alpha})_{(2)}^\intercal \boldsymbol{X}, \boldsymbol{X}] \\
&\quad + 4\operatorname{cov}[(\boldsymbol{I}_{K+1} \otimes \boldsymbol{Y}^\intercal)\mathcal{A}(\boldsymbol{\alpha})_{(2)}^\intercal \boldsymbol{X}, (\boldsymbol{I}_{K+1} \otimes \boldsymbol{Z}^\intercal)\mathcal{A}(\boldsymbol{\alpha})_{(2)}^\intercal \boldsymbol{X}] \quad \text{(S58)} \\
&= \operatorname{cov}[\boldsymbol{X}, \boldsymbol{X}] - 2\operatorname{cov}[\boldsymbol{X}, \boldsymbol{X}]\mathcal{A}(\boldsymbol{\alpha})_{(2)}(\boldsymbol{I}_{K+1} \otimes \boldsymbol{q}) - 2(\boldsymbol{I}_{K+1} \otimes \boldsymbol{q}^\intercal)\mathcal{A}(\boldsymbol{\alpha})_{(2)}^\intercal \operatorname{cov}[\boldsymbol{X}, \boldsymbol{X}] \\
&\quad + 4(\boldsymbol{I}_{K+1} \otimes \boldsymbol{q}^\intercal)\mathcal{A}(\boldsymbol{\alpha})_{(2)}^\intercal \operatorname{cov}[\boldsymbol{X}, \boldsymbol{X}]\mathcal{A}(\boldsymbol{\alpha})_{(2)}(\boldsymbol{I}_{K+1} \otimes \boldsymbol{q}) \quad \text{(S59)} \\
&= \boldsymbol{Q} - 2\boldsymbol{Q}\mathcal{A}(\boldsymbol{\alpha})_{(2)}(\boldsymbol{I}_{K+1} \otimes \boldsymbol{q}) - 2(\boldsymbol{I}_{K+1} \otimes \boldsymbol{q}^\intercal)\mathcal{A}(\boldsymbol{\alpha})_{(2)}^\intercal \boldsymbol{Q} \\
&\quad + 4(\boldsymbol{I}_{K+1} \otimes \boldsymbol{q}^\intercal)\mathcal{A}(\boldsymbol{\alpha})_{(2)}^\intercal \boldsymbol{Q}\mathcal{A}(\boldsymbol{\alpha})_{(2)}(\boldsymbol{I}_{K+1} \otimes \boldsymbol{q}) \quad \text{(S60)} \\
&= [\boldsymbol{I}_{K+1} - 2\mathcal{A}(\boldsymbol{\alpha})_{(2)}(\boldsymbol{I}_{K+1} \otimes \boldsymbol{q})]^\intercal \boldsymbol{Q}[\boldsymbol{I}_{K+1} - 2\mathcal{A}(\boldsymbol{\alpha})_{(2)}(\boldsymbol{I}_{K+1} \otimes \boldsymbol{q})], \quad \text{(S61)}
\end{aligned}
$$

where $\boldsymbol{Y}$ and $\boldsymbol{Z}$ are *iid* to $\boldsymbol{X}$. □

We will now develop a method to estimate $\boldsymbol{\alpha}$ given equilibrium, and to test for equilibrium while accounting for $\boldsymbol{\alpha}$. Since the expectation of the $U$-statistic is $\mathbf{0}$ under the null of equilibrium, our estimator of $\boldsymbol{\alpha}$, conditional on equilibrium, is the minimizer of the following quadratic form

$$
n\hat{V}_n(\boldsymbol{\alpha})^\intercal \boldsymbol{\Omega} \hat{V}_n(\boldsymbol{\alpha}), \quad \text{(S62)}
$$

where $\boldsymbol{\Omega}$ is a $(K+1) \times (K+1)$ symmetric positive semi-definite matrix. Consistency and asymptotic normality of the minimizer of (S62), say $\hat{\boldsymbol{\alpha}}$, is guaranteed by the theory on so-called "$U$-process minimizers" [Honoré and Powell, 1994, Bose, 2002, Bose and Chatterjee, 2018]. In order to come up with a test for equilibrium, we will connect the methods of $U$-process minimizers with those from the generalized methods-of-moments literature from econometrics [Hansen, 1982].

**Theorem S3.** *Let $\boldsymbol{\Omega} = \boldsymbol{\Sigma}^+$ be the Moore-Penrose inverse [Moore, 1920, Penrose, 1955] of $\boldsymbol{\Sigma}$, the asymptotic covariance of $\sqrt{n}\hat{V}_n(\hat{\boldsymbol{\alpha}})$. Then*

$$
n\hat{V}_n(\hat{\boldsymbol{\alpha}})^\intercal \boldsymbol{\Omega} \hat{V}_n(\hat{\boldsymbol{\alpha}}) \xrightarrow{\mathcal{L}} \chi^2_{\nu - \lfloor K/4 \rfloor} \quad \text{(S63)}
$$

*where $\nu$ is the rank of $\boldsymbol{\Omega}$.*

*Proof.* This is a special case of the Sargan-Hansen $J$-test [Hansen, 1982]. Let $\boldsymbol{\alpha}_0$ be the true value of $\boldsymbol{\alpha}$. It suffices to show that the following is an idempotent matrix of rank $\nu$.

$$
\begin{aligned}
\lim_{n\to\infty} \operatorname{cov}[\boldsymbol{\Omega}^{1/2}\hat{V}_n(\boldsymbol{\alpha}_0)] &= \boldsymbol{\Omega}^{1/2} \lim_{n\to\infty} \operatorname{cov}[\hat{V}_n(\boldsymbol{\alpha}_0)]\boldsymbol{\Omega}^{1/2} \quad \text{(S64)} \\
&= \boldsymbol{\Omega}^{1/2}\boldsymbol{\Sigma}\boldsymbol{\Omega}^{1/2} \quad \text{(S65)} \\
&= \boldsymbol{\Sigma}^{+1/2}\boldsymbol{\Sigma}\boldsymbol{\Sigma}^{+1/2} \quad \text{(S66)} \\
& \quad \text{(S67)}
\end{aligned}
$$

One way to represent the Moore-Penrose inverse is in terms of its eigenvalue decomposition:

$$
\begin{aligned}
\boldsymbol{\Sigma} &= \boldsymbol{U}\boldsymbol{D}\boldsymbol{U}^\intercal \quad \text{(S68)} \\
\boldsymbol{\Sigma}^+ &= \boldsymbol{U}\boldsymbol{F}\boldsymbol{U}^\intercal, \quad \text{(S69)}
\end{aligned}
$$

where $\boldsymbol{F} = \text{diag}(1/D_{11}, \dots, 1/D_{\nu\nu}, 0, \dots, 0)$. Thus

$$(\text{S66}) = \boldsymbol{U}\boldsymbol{F}^{1/2}\boldsymbol{U}^\intercal\boldsymbol{U}\boldsymbol{D}\boldsymbol{U}^\intercal\boldsymbol{U}\boldsymbol{F}^{1/2}\boldsymbol{U}^\intercal = \boldsymbol{U}\boldsymbol{I}\boldsymbol{U}^\intercal, \tag{S70}$$

where $\boldsymbol{I} = \text{diag}(1, \dots, 1, 0, \dots, 0)$, having $K + 1 - \nu$ zeros and $\nu$ non-zeros. This is an idempotent matrix of rank $\nu$. □

The result in Theorem S3 is for when we know $\boldsymbol{\Sigma}$, but using Slutsky's theorem, it suffices to use a consistent estimate of $\boldsymbol{\Sigma}$. This requires knowledge of $\boldsymbol{\alpha}$. Our solution is to use a multistep approach, which we will describe now. In the first step, we obtain a consistent estimate of $\boldsymbol{\alpha}$ by using $\boldsymbol{\Omega} = \boldsymbol{I}_{K+1}$. Call this consistent estimate $\tilde{\boldsymbol{\alpha}}$. We then use this estimate to obtain a consistent estimate of $\boldsymbol{\Sigma}$:

$$\hat{\boldsymbol{\Sigma}} := [\boldsymbol{I}_{K+1} - 2\mathcal{A}(\tilde{\boldsymbol{\alpha}})_{(2)}(\boldsymbol{I}_{K+1} \otimes \hat{\boldsymbol{f}})]^\intercal[\text{diag}(\hat{\boldsymbol{f}}) - \hat{\boldsymbol{f}}\hat{\boldsymbol{f}}^\intercal][\boldsymbol{I}_{K+1} - 2\mathcal{A}(\tilde{\boldsymbol{\alpha}})_{(2)}(\boldsymbol{I}_{K+1} \otimes \hat{\boldsymbol{f}})], \text{ where} \tag{S71}$$

$$\hat{\boldsymbol{f}} := f(\hat{\boldsymbol{q}}, \tilde{\boldsymbol{\alpha}}). \tag{S72}$$

We use $\hat{\boldsymbol{f}}$ instead of $\hat{\boldsymbol{q}}$ in (S71) because, under the null $\boldsymbol{q} = f(\boldsymbol{q}, \boldsymbol{\alpha})$, and so this covariance is only valid given equilibrium, which should increase power of the test. Using $\hat{\boldsymbol{f}}$ also results in a direct generalization of the classical $\chi^2$ test from diploids to polyploids, whereas using $\hat{\boldsymbol{q}}$ would not (Section S8). We finally use $\boldsymbol{\Omega} = \hat{\boldsymbol{\Sigma}}^+$ to obtain our final estimate of $\boldsymbol{\alpha}$, called $\hat{\boldsymbol{\alpha}}$, and run a hypothesis test for equilibrium. A summary of this approach is described in Procedure 1.

##### S7.1 Notes on $U$-statistic approach

**Note 1:** Any positive semi-definite weight matrix $\boldsymbol{\Omega}$ used in (S62) will yield a consistent estimate of $\boldsymbol{\alpha}$. But only setting $\boldsymbol{\Omega}$ to be a generalized inverse of $\boldsymbol{\Sigma}$ will result in a proper $\chi^2$ distribution for the test-statistic. This motivates the two-step procedure. We can obtain a consistent estimate of $\boldsymbol{\alpha}$, use this to obtain a consistent estimate of $\boldsymbol{\Sigma}$, which guarantees the asymptotic distribution of the test-statistic.

**Note 2:** The naive way to calculate the $U$- and $V$-statistics (Equations (S40) and (S42)) has computational complexity of $\mathcal{O}(n^3)$. However, because of Theorems S1 and S2, none of the operations in are actually $\mathcal{O}(n^3)$. But rather they all have complexity $\mathcal{O}(n)$, the cost to calculate $\hat{\boldsymbol{q}}$.

**Note 3:** Numerical results indicate that the covariance (S53) has rank $\nu = K - 1$, so two less than full rank. However, for robustness, we estimate the rank using numerical methods each time we run Procedure 1.

**Note 4:** Though we have asymptotic results for the distribution of $T(\hat{\boldsymbol{\alpha}})$, the approximations can be really bad for small $n$ because there are very few individuals with some genotypes. However, we can improve the approximation (at the risk of losing power) by combining genotypes together. We will describe this procedure now. We have $\sqrt{n}\hat{V}_n(\boldsymbol{\alpha}) \xrightarrow{\mathcal{L}} N(0, \boldsymbol{\Sigma})$. Without loss of generality, let us suppose that we are combining the last $m$ variables. Then this transformation can be represented by $\boldsymbol{H}\hat{V}_n(\boldsymbol{\alpha})$, where

$$\boldsymbol{H} = \begin{pmatrix} \boldsymbol{I}_{K+1-m} & \boldsymbol{0}_{(K+1-m)\times m} \\ \boldsymbol{0}_{K+1-m}^\intercal & \boldsymbol{1}_m^\intercal \end{pmatrix}. \tag{S73}$$

Thus, using the $\delta$-method, we have

$$\sqrt{n}\boldsymbol{H}\hat{V}_n(\boldsymbol{\alpha}) \xrightarrow{\mathcal{L}} N(0, \boldsymbol{H}\boldsymbol{\Sigma}\boldsymbol{H}^\intercal). \tag{S74}$$

Then we have

$$n\hat{V}_n(\boldsymbol{\alpha})^\intercal \boldsymbol{H}^\intercal[\boldsymbol{H}\boldsymbol{\Sigma}\boldsymbol{H}^\intercal]^+\boldsymbol{H}\hat{V}_n(\boldsymbol{\alpha}) \xrightarrow{\mathcal{L}} \chi^2_{\nu-\lfloor K/4\rfloor}, \tag{S75}$$

where $\nu$ is the rank of $\boldsymbol{H}\boldsymbol{\Sigma}\boldsymbol{H}^\intercal$, which we find numerically.

By default, we pool all genotypes that have zero individuals. If pooling results in fewer groups than $\lfloor K/4\rfloor + 1$, then we do not run the test.

**Note 5:** It is well-known that the rates of double reduction are bounded under certain models of meiosis [Huang et al., 2019]. To use this information, when we optimize over $\boldsymbol{\alpha}$ to obtain either $\tilde{\boldsymbol{\alpha}}$ or $\hat{\boldsymbol{\alpha}}$, we do so under the constraint $0 \leq \alpha_i \leq c_i$ for a theoretical maximum value of $c_i$. These maximum values come from the complete equational segregation model, as derived in Huang et al. [2019] (their equations (4) and (5)). This is the model that provides the largest bounds on the rates of double reduction in the literature. Specifically, $\alpha_i$ in a $K$-ploid species is allowed to vary by

$$0 \leq \alpha_i \leq \sum_{j=i}^{\lfloor K/4\rfloor} 2^{K/2-3j} \frac{\binom{j}{i}\binom{K/2}{j}\binom{K/2-j}{K/2-2j}}{\binom{K}{K/2}}. \tag{S76}$$

**Note 6:** One issue that arises when we optimize over $0 \leq \alpha_i \leq c_i$ from (S76) is that the asymptotic result of Theorem S3 no longer holds if $\alpha_i$ is exactly 0 or $c_i$, because the asymptotics require the parameters to be at an interior point in the parameterspace. Also, for $\alpha_i$ close to either 0 or $c_i$, the asymptotics can be really bad and require very large samples to be an adequate approximation. One *ad-hoc* workaround that we found works really well in practice is to increase the degrees of freedom of the $\chi^2$ distribution by the number of $\hat{\alpha}_i$'s estimated at the boundary. The intuition is that if $\hat{\alpha}_i$ is fixed and on the boundary, then we should increase the degrees of freedom. This is the procedure we used in the simulations of Section 3.1.

### S8 Diploid procedure

In this section, we prove that in diploids, Procedure 1 is equivalent to the classical $\chi^2$-test for HWE. Note that in diploids, there is no $\boldsymbol{\alpha}$, and so this results in a much simplified procedure.

In diploids, the genotype frequencies are denoted $\boldsymbol{q} = (q_0, q_1, q_2)^\intercal$. Since there are two degrees of freedom for genotype frequencies, we may represent these genotype frequencies in terms of two parameters, the allele frequency $r$, and the disequilibrium coefficient $t$.

$$r = \frac{1}{2}q_1 + q_2, \tag{S77}$$

$$t = q_2 - r^2, \tag{S78}$$

which implies that

$$q_0 = (1-r)^2 + t, \tag{S79}$$
$$q_1 = 2r(1-r) - 2t, \tag{S80}$$
$$q_2 = r^2 + t. \tag{S81}$$

The classical $\chi^2$ procedure, using estimated genotype frequencies $\hat{\boldsymbol{q}} = (\hat{q}_0, \hat{q}_1, \hat{q}_2)^\intercal$ and sample size $n$, is [Weir, 1996]

$$\hat{r} = \frac{1}{2}\hat{q}_1 + \hat{q}_2, \tag{S82}$$
$$\hat{t} = \hat{q}_2 - \hat{r}^2, \tag{S83}$$
$$\frac{n\hat{t}^2}{\hat{r}^2(1-\hat{r})^2} \sim \chi_1^2. \tag{S84}$$

The following contains the genotype frequencies after one round of random mating in diploids in terms of the previous generation's genotype frequencies. However, since HWE is reached after one generation of random mating in diploids, one can easily show that these are equal to the HWE frequencies:

$$f(\boldsymbol{q}) = \begin{pmatrix} q_0^2 + q_0 q_1 + \frac{1}{4}q_1^2 \\ q_0 q_1 + 2q_0 q_2 + \frac{1}{2}q_1^2 + q_1 q_2 \\ \frac{1}{4}q_1^2 + q_1 q_2 + q_2^2 \end{pmatrix} = \begin{pmatrix} (1-r)^2 \\ 2r(1-r) \\ r^2 \end{pmatrix}, \tag{S85}$$

and so

$$\boldsymbol{q} - f(\boldsymbol{q}) = \begin{pmatrix} q_0 - (1-r)^2 \\ q_1 - 2r(1-r) \\ q_2 - r^2 \end{pmatrix} = \begin{pmatrix} t \\ -2t \\ t \end{pmatrix}, \tag{S86}$$

which means that $\hat{\boldsymbol{q}} - f(\hat{\boldsymbol{q}}) = (\hat{t}, -2\hat{t}, \hat{t})^\intercal$.

To obtain the asymptotic covariance of $\hat{\boldsymbol{q}} - f(\hat{\boldsymbol{q}})$, we need the $\boldsymbol{A}$ array, which in diploids is

$$\mathcal{A} = \left(\begin{array}{ccc|ccc|ccc} 1 & 1/2 & 0 & 1/2 & 1/4 & 0 & 0 & 0 & 0 \\ 0 & 1/2 & 1 & 1/2 & 1/2 & 1/2 & 1 & 1/2 & 0 \\ 0 & 0 & 0 & 0 & 1/4 & 1/2 & 0 & 1/2 & 1 \end{array}\right), \tag{S87}$$

where the vertical lines differentiate the third index. We have

$$\mathcal{A}(\boldsymbol{\alpha})_{(2)}(\boldsymbol{I}_{K+1} \otimes \boldsymbol{f}) = \begin{pmatrix} f_0 + \frac{1}{2}f_1 & \frac{1}{2}f_1 + f_2 & 0 \\ \frac{1}{2}f_0 + \frac{1}{4}f_1 & \frac{1}{2}f_0 + \frac{1}{2}f_1 + \frac{1}{2}f_2 & \frac{1}{4}f_1 + \frac{1}{2}f_2 \\ 0 & f_0 + \frac{1}{2}f_1 & \frac{1}{2}f_1 + f_2 \end{pmatrix}. \tag{S88}$$

We can use this to obtain an estimate of the covariance matrix (S53). Using Mathematica [Wolfram Research, Inc., 2020] to simplify expressions, we have that (see online material `https://github.`

com/dcgerard/hwesims)

$$\hat{\mathbf{\Sigma}} = \hat{r}^2(1-\hat{r})^2 \begin{pmatrix} 1 & -2 & 1 \\ -2 & 4 & -2 \\ 1 & -2 & 1 \end{pmatrix} \tag{S89}$$

$$= 6\hat{r}^2(1-\hat{r})^2\boldsymbol{u}_1\boldsymbol{u}_1^2, \tag{S90}$$

where $\boldsymbol{u}_1 = (1, -2, 1)/\sqrt{6}$. Equation (S90) is the eigenvalue decomposition of $\hat{\mathbf{\Sigma}}$ with first eigenvalue $d_1 = 6\hat{r}^2(1-\hat{r})^2$ and first eigenvector $\boldsymbol{u}_1$. This means that

$$\mathbf{\Sigma}^+ = \frac{1}{d_1}\boldsymbol{u}_1\boldsymbol{u}_1^\intercal. \tag{S91}$$

Our test-statistic is

$$n[\hat{\boldsymbol{q}} - f(\hat{\boldsymbol{q}})]^\intercal\mathbf{\Sigma}^+[\hat{\boldsymbol{q}} - f(\hat{\boldsymbol{q}})] = \frac{n\{[\hat{\boldsymbol{q}} - f(\hat{\boldsymbol{q}})]^\intercal\boldsymbol{u}_1\}^2}{\hat{d}_1} \tag{S92}$$

$$= \frac{n[(\hat{t} + 4\hat{t} + \hat{t})/\sqrt{6}]^2}{6\hat{r}^2(1-\hat{r})^2} \tag{S93}$$

$$= \frac{n\hat{t}^2}{\hat{r}^2(1-\hat{r})^2}, \tag{S94}$$

which we compare to a $\chi_1^2$ distribution. Comparing (S84) to (S94), we see that the $U$-statistic approach is equivalent to the classical $\chi^2$-test in diploids.

One can show that if we would have used $\hat{\boldsymbol{q}}$ instead of $\hat{\boldsymbol{f}}$ to obtain $\hat{\mathbf{\Sigma}}$, we would have obtained $d_1 = 6(r^2(1-r)^2 + (1-2r)^2t - t^2)$ (see online material https://github.com/dcgerard/hwesims), and the resulting variance estimate in the denominator of (S94) would have been that under the alternative, rather than the null (see equation (3.3) of Weir [1996]). This motivates using using $\hat{\boldsymbol{f}}$ in (S71) as the using the null variance is the more principled approach, but both ways would asymptotically control for Type I error.

#### S9 Derivative of $f(\boldsymbol{q}, \boldsymbol{\alpha})$ with respect to $\boldsymbol{\alpha}$

In this section, we will derive the Jacobian of (S27) with respect to $\boldsymbol{\alpha}$. This is used for the gradient calculations during the gradient ascent during the minimization of (S62). For convenience, we will drop the dependence on $\boldsymbol{q}$. Thus, we have

$$f(\boldsymbol{\alpha}) = \boldsymbol{g} \circ \boldsymbol{p} \circ \boldsymbol{B}(\boldsymbol{\alpha}), \tag{S95}$$

where

$$B(\boldsymbol{\alpha})_{\ell k} = \sum_{i=0}^{\lfloor K/4 \rfloor} \sum_{j=0}^{i} \frac{\binom{\ell}{j}\binom{\ell-j}{k-2j}\binom{K-\ell}{i-j}\binom{K-\ell-(i-j)}{K/2-k-2(i-j)}}{\binom{K}{i}\binom{K-i}{K/2-2i}}\alpha_i, \tag{S96}$$

$$\boldsymbol{p}(\boldsymbol{B}) = \boldsymbol{B}^\intercal\boldsymbol{q}, \text{ and} \tag{S97}$$

$$\boldsymbol{g}(\boldsymbol{p}) = \boldsymbol{p} * \boldsymbol{p}. \tag{S98}$$

We can therefore use the chain rule to obtain the Jacobian $\mathrm{D}f(\boldsymbol{\alpha})$:

$$\mathrm{D}f(\boldsymbol{\alpha}) = (\mathrm{D}\boldsymbol{g}(\boldsymbol{p}))(\mathrm{D}\boldsymbol{p}(\boldsymbol{B}))(\mathrm{D}\boldsymbol{B}(\boldsymbol{\alpha})). \tag{S99}$$

One can verify that $\mathrm{D}\boldsymbol{g}(\boldsymbol{p})$ is the following $(K+1)\times(K/2+1)$ Toeplitz matrix

$$\mathrm{D}\boldsymbol{g}(\boldsymbol{p}) = \begin{pmatrix} p_0 & 0 & & & & \\ p_1 & p_0 & 0 & & & \mathbf{0} \\ p_2 & p_1 & \ddots & \ddots & & \\ \vdots & p_2 & \ddots & \ddots & \ddots & \\ p_{K/2-1} & \vdots & \ddots & \ddots & \ddots & 0 \\ p_{K/2} & p_{K/2-1} & & \ddots & \ddots & p_0 \\ 0 & p_{K/2} & \ddots & & \ddots & p_1 \\ & 0 & \ddots & \ddots & & p_2 \\ & & \ddots & \ddots & \ddots & \vdots \\ \mathbf{0} & & & \ddots & \ddots & p_{K/2-1} \\ & & & & 0 & p_{K/2} \end{pmatrix} \tag{S100}$$

One can also verify that $\mathrm{D}\boldsymbol{p}(\boldsymbol{B})$ is the following $(K/2+1)\times(K+1)(K/2+1)$ matrix:

$$\mathrm{D}\boldsymbol{p}(\boldsymbol{B}) = \boldsymbol{I}_{K/2+1} \otimes \boldsymbol{q}^{\intercal}, \tag{S101}$$

where "$\otimes$" is the Kronecker product and $\boldsymbol{I}_{K/2+1}$ is the identity matrix in $K/2+1$ dimensions. Finally, because (S96) is linear in the $\alpha_i$'s, we have that $\mathrm{D}\boldsymbol{B}(\boldsymbol{\alpha})$ is a $(K+1)(K/2+1)\times\lfloor K/4\rfloor$ matrix (setting $\alpha_0 = 1-\sum_{i=1}^{\lfloor K/4\rfloor}\alpha_i$ and only taking derivatives with for $\alpha_i$ with $i>0$) with elements:

$$(\mathrm{D}\boldsymbol{B}(\boldsymbol{\alpha}))_{r(\ell,k),i} = \sum_{j=0}^{i} \frac{\binom{\ell}{j}\binom{\ell-j}{k-2j}\binom{K-\ell}{i-j}\binom{K-\ell-(i-j)}{K/2-k-2(i-j)}}{\binom{K}{i}\binom{K-i}{K/2-2i}} - \frac{\binom{\ell}{k}\binom{K-\ell}{K/2-k}}{\binom{K}{K/2}}, \tag{S102}$$

where $r(\ell,k)$ is a function that maps the $(\ell,k)$ index of a matrix to the $r(\ell,k)$ index of the vectorization of that matrix. Specifically:

$$r(\ell,k) = 1+\ell+(K+1)k. \tag{S103}$$

The only term in the Jacobian in (S99) that changes from iteration to iteration during a gradient ascent is $\mathrm{D}\boldsymbol{g}(\boldsymbol{p})$, and so $\mathrm{D}\boldsymbol{p}(\boldsymbol{B})\mathrm{D}\boldsymbol{B}(\boldsymbol{\alpha})$ may be pre-calculated to improve computational performance.

#### S10 A likelihood ratio test given no double reduction

In the absence of double reduction, we can test for equilibrium, specifically HWE. Under the null of equilibrium, we have MLE genotype frequencies of $\hat{\boldsymbol{q}}_0$ from (15). Under the alternative of violations in equilibrium, we have MLE genotype frequencies of $\hat{\boldsymbol{q}} = \boldsymbol{y}/n$. The following likelihood ratio test statistic follows a $\chi^2_{K-1}$ distribution under the null

$$-2[\log \text{Multinom}(\boldsymbol{y}|n, \hat{\boldsymbol{q}}_0) - \log \text{Multinom}(\boldsymbol{y}|n, \hat{\boldsymbol{q}})], \tag{S104}$$

which we can use to obtain a $p$-value for this test.

#### S11 An initial exploration on the effects of genotype uncertainty

We ran a small simulation study on the effects of genotype uncertainty on our equilibrium testing procedures. We generated genotypes from $n = 1000$ individuals of ploidy $K = 6$ after either $m = 2$ or $m = \infty$ rounds of random mating (using Procedure 3). Equilibrium is fulfilled at $m = \infty$ and is not fulfilled at $m = 2$, but random mating is fulfilled in both scenarios. We used an allele frequency of $r = 0.5$ and a rate of double reduction of $\alpha = 0.15$, which is half the maximum rate for hexaploids under the complete equational segregation model (S76). We then used `rflexdog()` from `updog` package [Gerard et al., 2018, Gerard and Ferrão, 2019] to generate read counts (of depth 30) with overdispersion parameter 0.01, no allele bias, and a sequencing error rate of 0.01. We then used `updog` to obtain genotype posterior probabilities using these read counts. We used these probabilities directly in our bootstrap procedure (Section 2.5) to test for equilibrium. For the other procedures, we used the posterior expectation for the number of each genotype. We ran 100 replications for each scenario.

Histograms of the $p$-values are presented in Figure S28. We see there that the equilibrium testing procedures are all conservative when equilibrium is satisfied (except for the method of Section S10 that does not account for double reduction). They are all powerful when equilibrium is not satisfied. The test for random mating is conservative at equilibrium, but slightly anti-conservative not at equilibrium, even though random mating is fulfilled in both scenarios.

#### S12 Supplementary figures

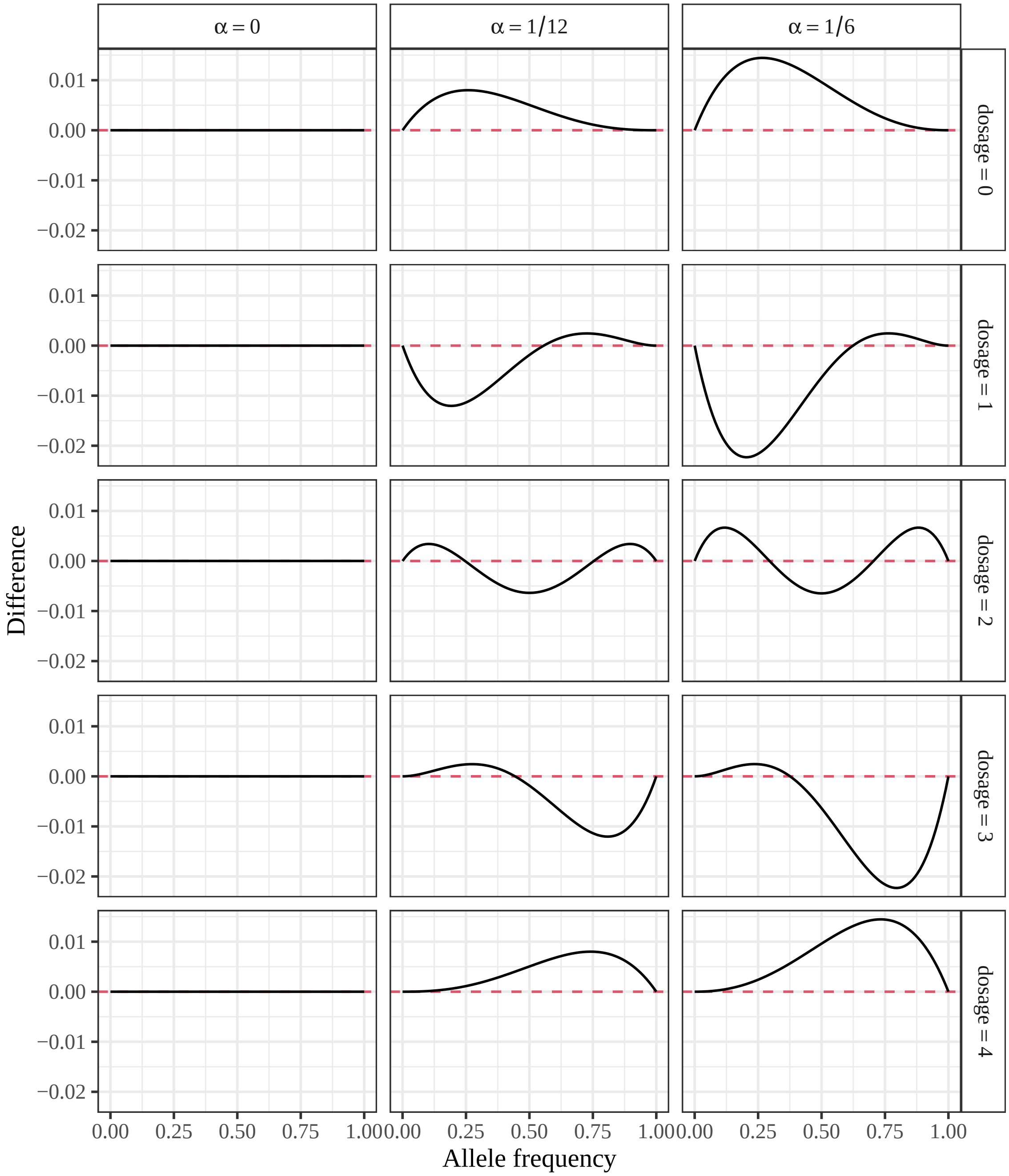


Figure S1: True allele frequency ($x$-axis) plotted against the difference in genotype frequencies calculated using the theoretical approach (Section 2.1) and that of Jiang et al. [2021] for each dosage level (row facets) and different levels of double reduction (column facets). Curves above zero (horizontal dashed line) indicate that the theoretical frequencies are larger than those calculated in Jiang et al. [2021].

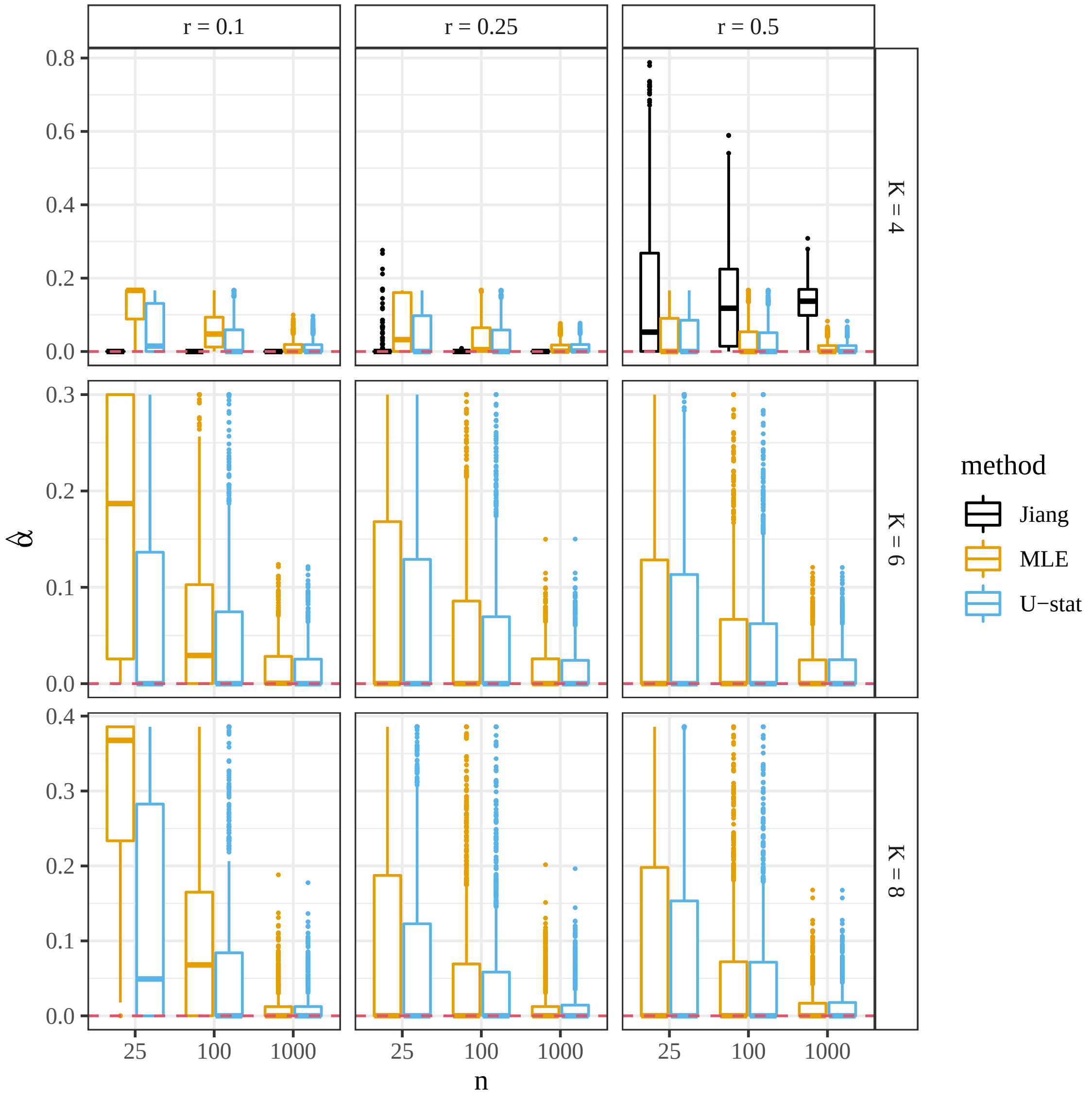


Figure S2: Estimates of $\alpha_1$ ($y$-axis) stratified by sample size ($x$-axis), ploidy (row facets), allele frequency (column facets), and method (color). The orange method is the MLE (Section 2.3), the blue is the $U$-statistic approach (Section 2.4), and the black is the method of [Jiang et al., 2021]. The true $\alpha_1$ is 0, also represented by the horizontal dashed line, and boxplots close to that line perform well.

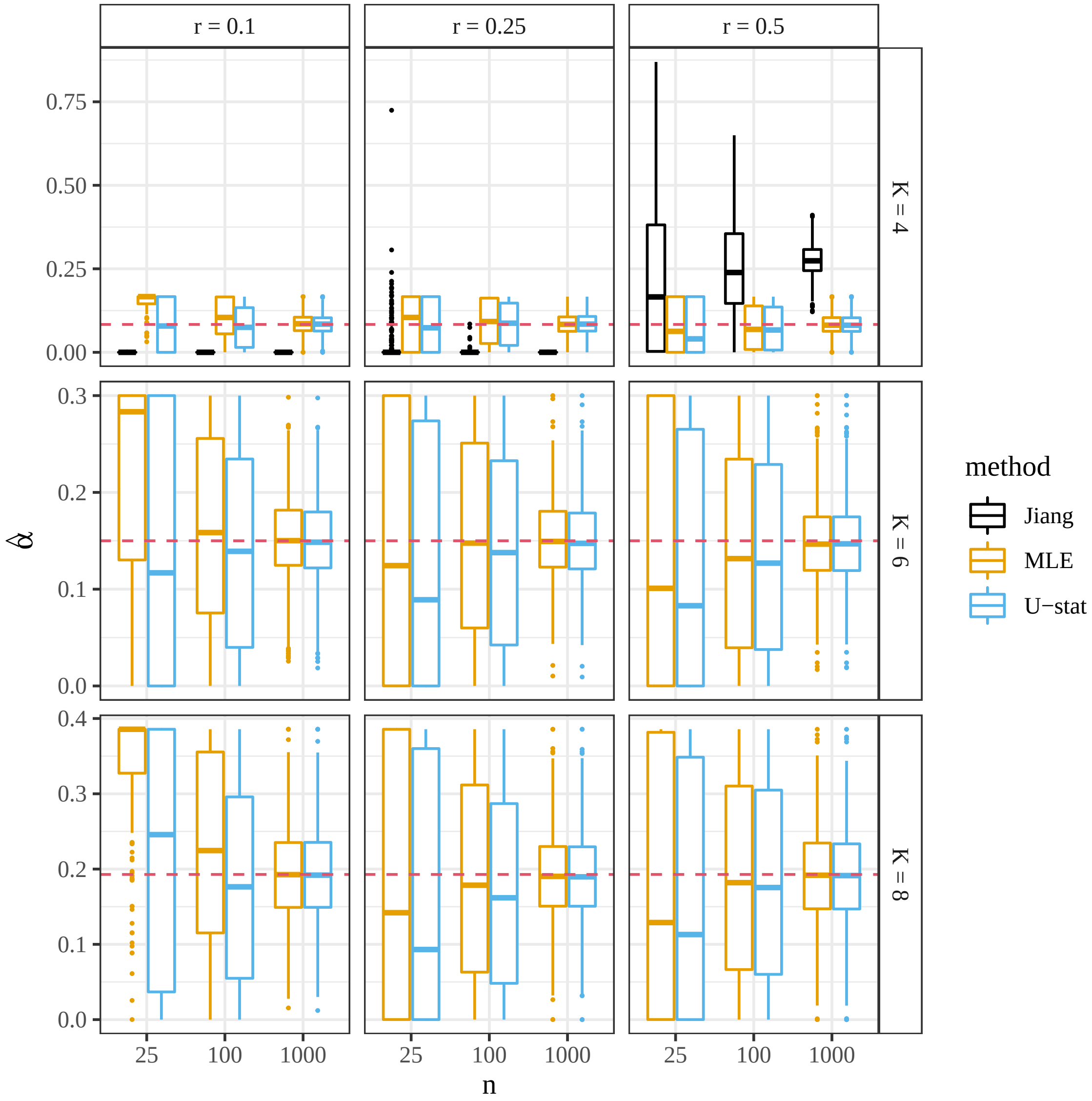


Figure S3: Estimates of $\alpha_1$ ($y$-axis) stratified by sample size ($x$-axis), ploidy (row facets), allele frequency (column facets), and method (color). The orange method is the MLE (Section 2.3), the blue is the $U$-statistic approach (Section 2.4), and the black is the method of [Jiang et al., 2021]. The true $\alpha_1$ is $\alpha_{1m}/2$, where $\alpha_{1m}$ is the maximum possible double reduction rate under the complete equational segregation model (Section S7.1). $\alpha_{1m}/2$ is also represented by the horizontal dashed line, and boxplots close to that line perform well.

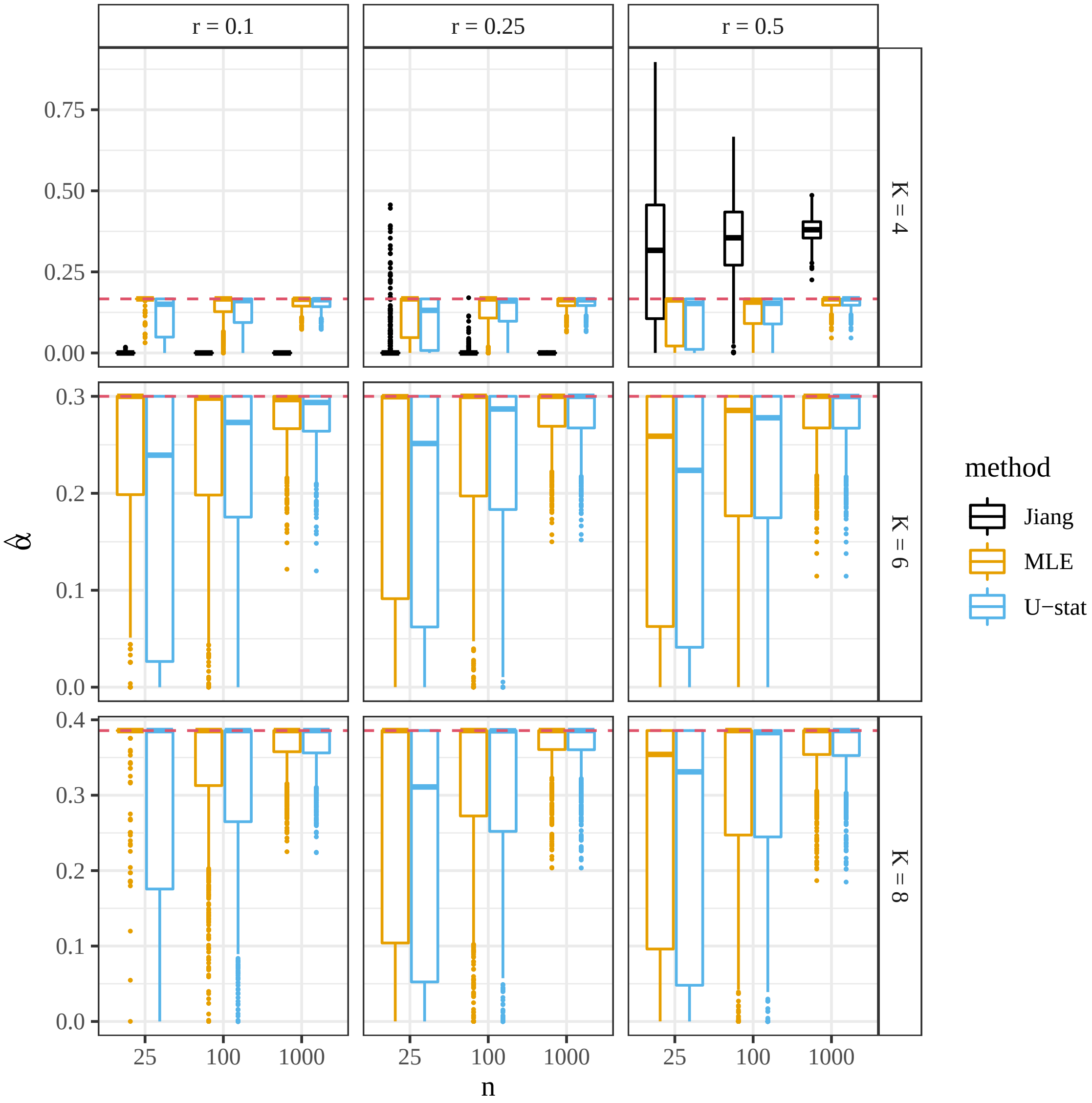


Figure S4: Estimates of $\alpha_1$ ($y$-axis) stratified by sample size ($x$-axis), ploidy (row facets), allele frequency (column facets), and method (color). The orange method is the MLE (Section 2.3), the blue is the $U$-statistic approach (Section 2.4), and the black is the method of [Jiang et al., 2021]. The true $\alpha_1$ is $\alpha_{1m}$, the maximum possible double reduction rate under the complete equational segregation model (Section S7.1). $\alpha_{1m}$ is also represented by the horizontal dashed line, and boxplots close to that line perform well.

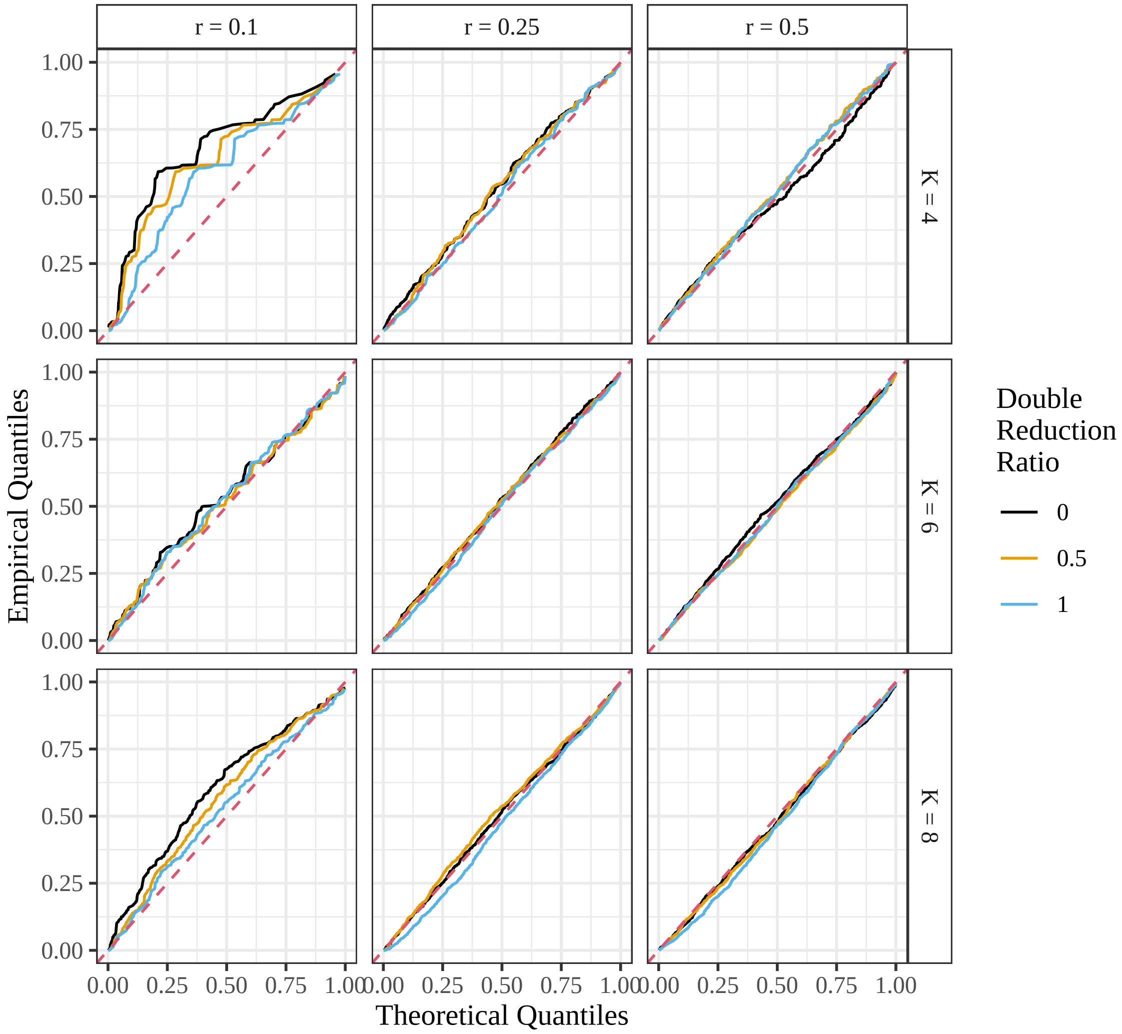


Figure S5: Quantile-quantile plots of the $U$-statistic approach's $p$-values (Section 2.4), testing against the null of equilibrium, when the sample size is $n = 25$, stratified by ploidy (row facets), allele frequency (column facets), and ratio of the double reduction parameter to the maximum possible double reduction rate from Section S7.1 (color). These $p$-values were calculated in settings where the null was satisfied, and so should lie on the $y = x$ line (red-dashed line).

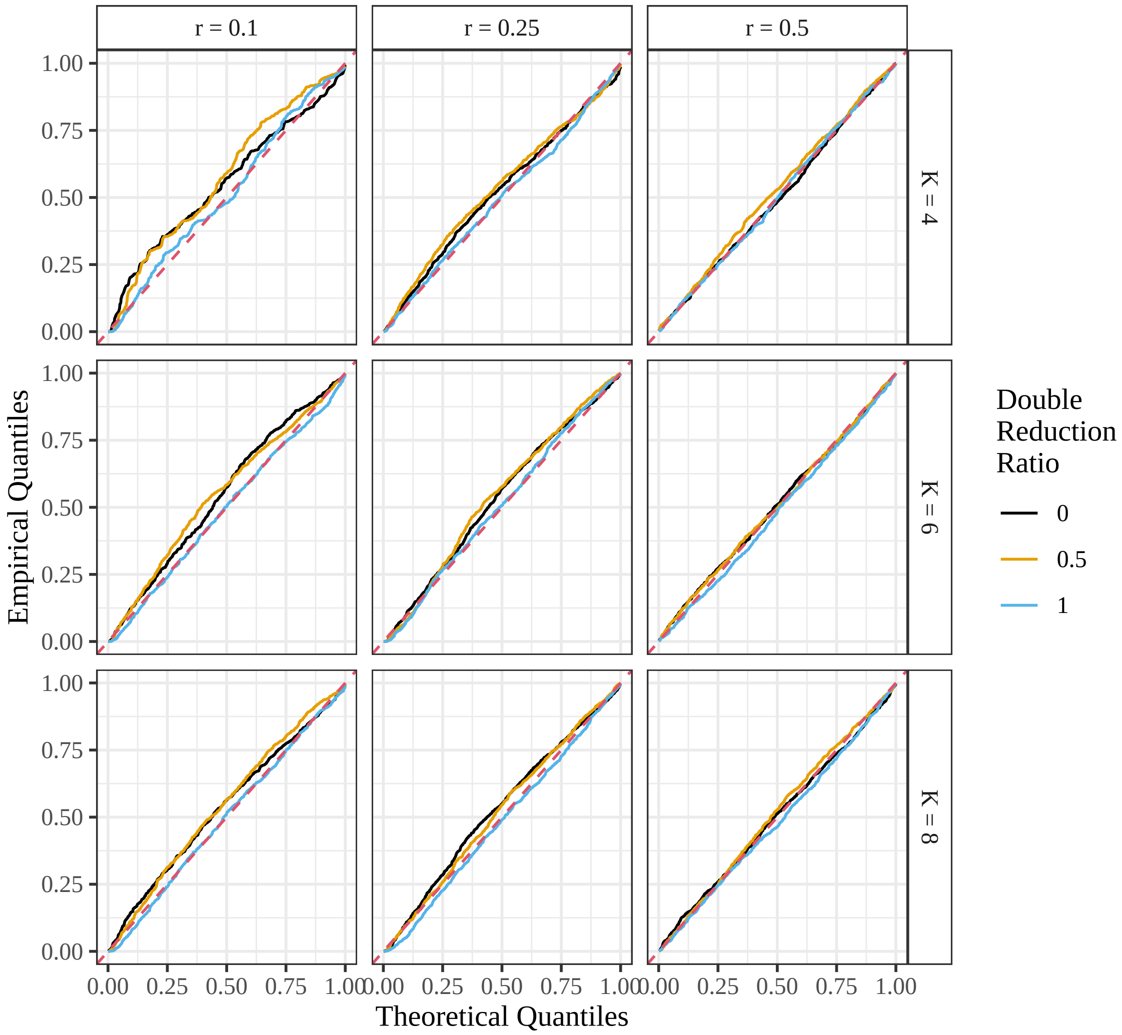


Figure S6: Quantile-quantile plots of the $U$-statistic approach's $p$-values (Section 2.4), testing against the null of equilibrium, when the sample size is $n = 100$, stratified by ploidy (row facets), allele frequency (column facets), and ratio of the double reduction parameter to the maximum possible double reduction rate from Section S7.1 (color). These $p$-values were calculated in settings where the null was satisfied, and so should lie on the $y = x$ line (red-dashed line).

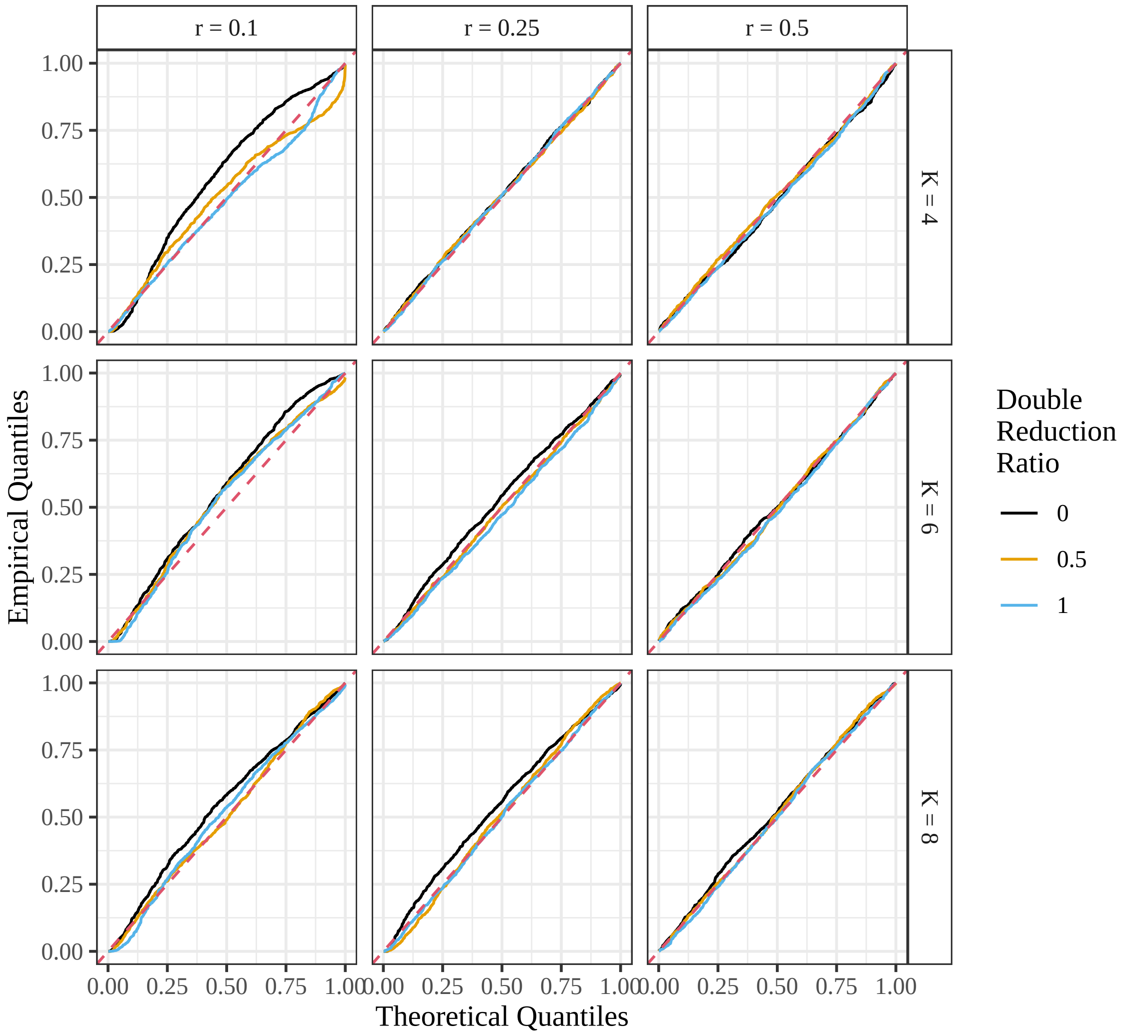


Figure S7: Quantile-quantile plots of the $U$-statistic approach's $p$-values (Section 2.4), testing against the null of equilibrium, when the sample size is $n = 1000$, stratified by ploidy (row facets), allele frequency (column facets), and ratio of the double reduction parameter to the maximum possible double reduction rate from Section S7.1 (color). These $p$-values were calculated in settings where the null was satisfied, and so should lie on the $y = x$ line (red-dashed line).

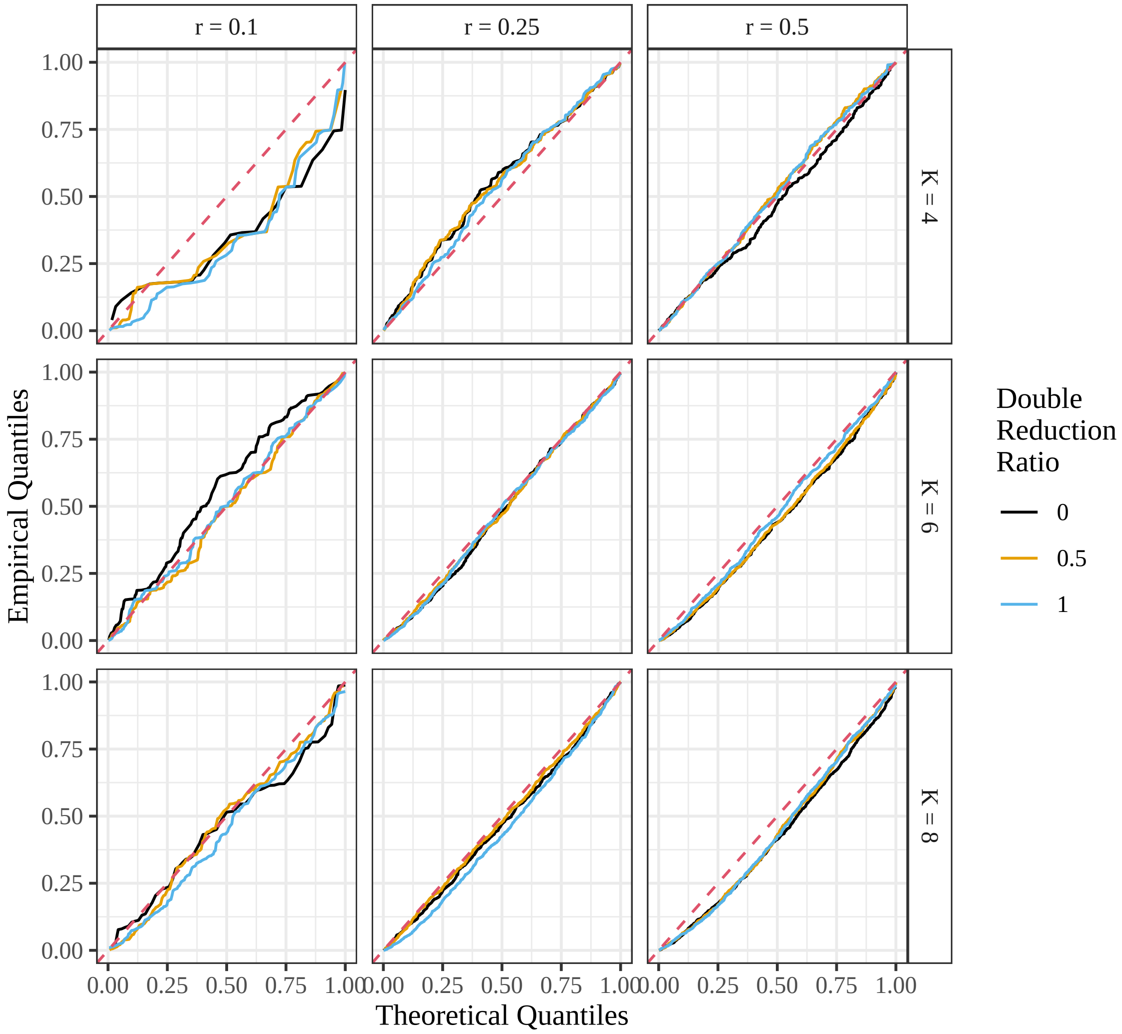


Figure S8: Quantile-quantile plots of the likelihood ratio test's approach's $p$-values (Section 2.3), testing against the null of equilibrium, when the sample size is $n = 25$, stratified by ploidy (row facets), allele frequency (column facets), and ratio of the double reduction parameter to the maximum possible double reduction rate from Section S7.1 (color). These $p$-values were calculated in settings where the null was satisfied, and so should lie on the $y = x$ line (red-dashed line).

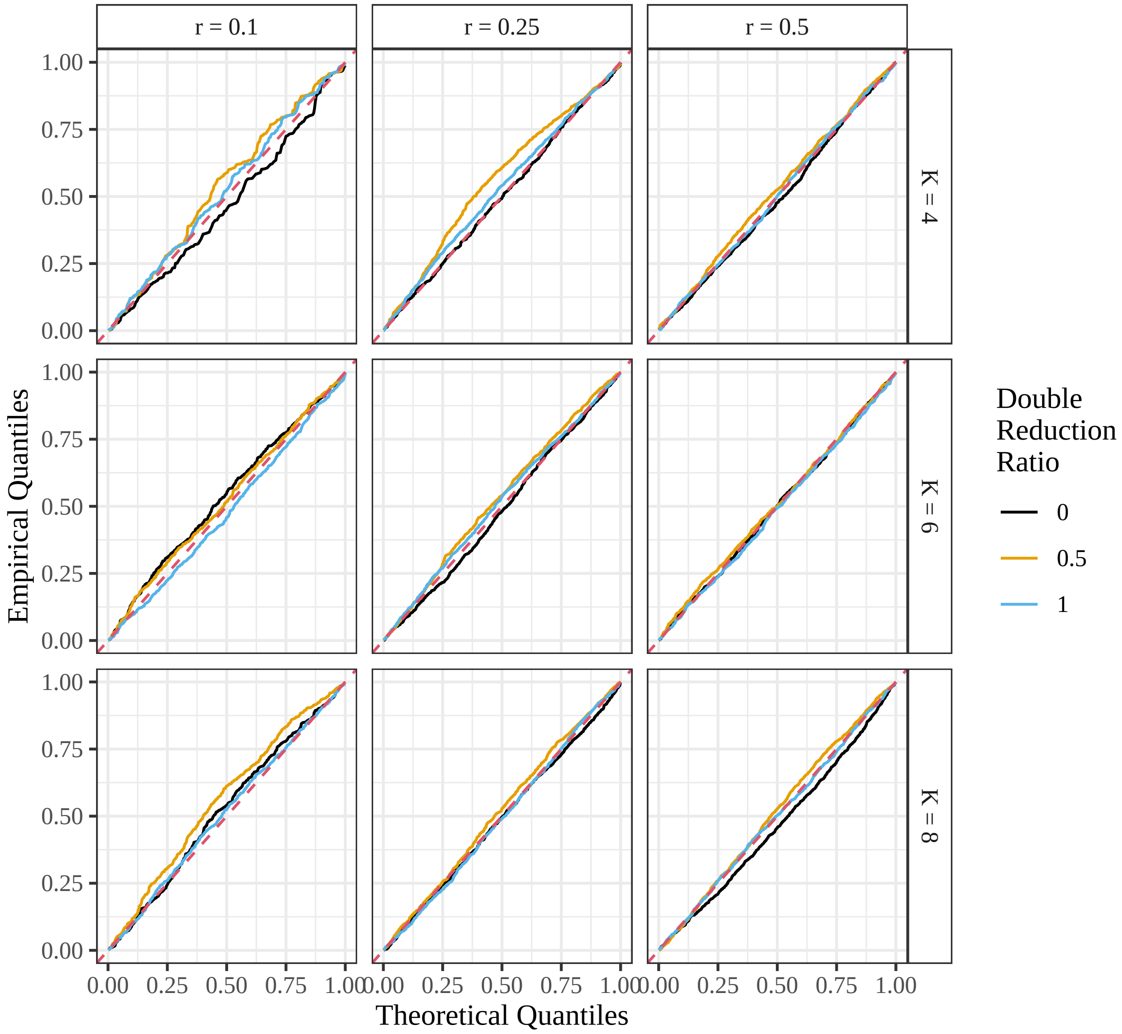


Figure S9: Quantile-quantile plots of the likelihood ratio test's approach's $p$-values (Section 2.3), testing against the null of equilibrium, when the sample size is $n = 100$, stratified by ploidy (row facets), allele frequency (column facets), and ratio of the double reduction parameter to the maximum possible double reduction rate from Section S7.1 (color). These $p$-values were calculated in settings where the null was satisfied, and so should lie on the $y = x$ line (red-dashed line).

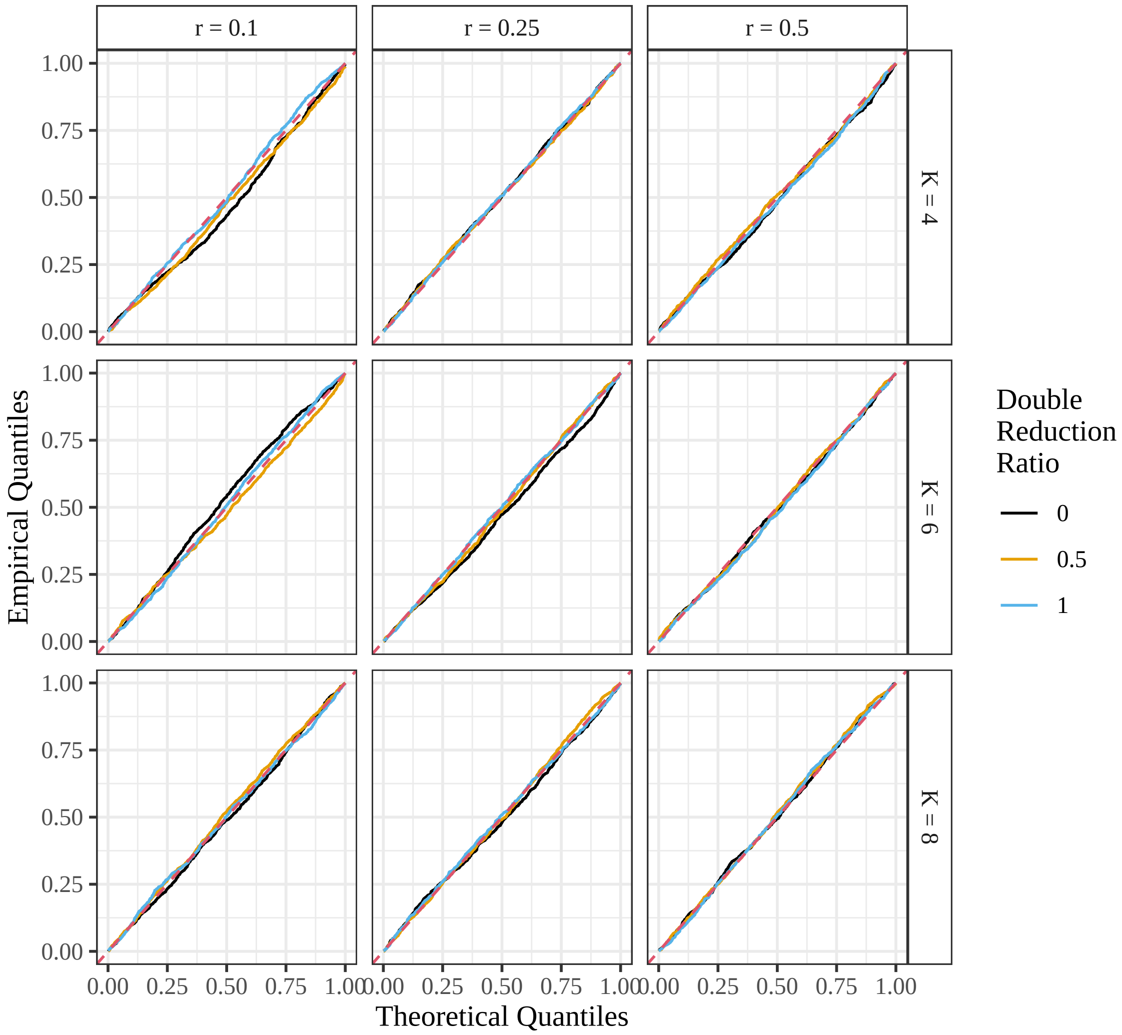


Figure S10: Quantile-quantile plots of the likelihood ratio test's $p$-values (Section 2.3), testing against the null of equilibrium, when the sample size is $n = 1000$, stratified by ploidy (row facets), allele frequency (column facets), and ratio of the double reduction parameter to the maximum possible double reduction rate from Section S7.1 (color). These $p$-values were calculated in settings where the null was satisfied, and so should lie on the $y = x$ line (red-dashed line).

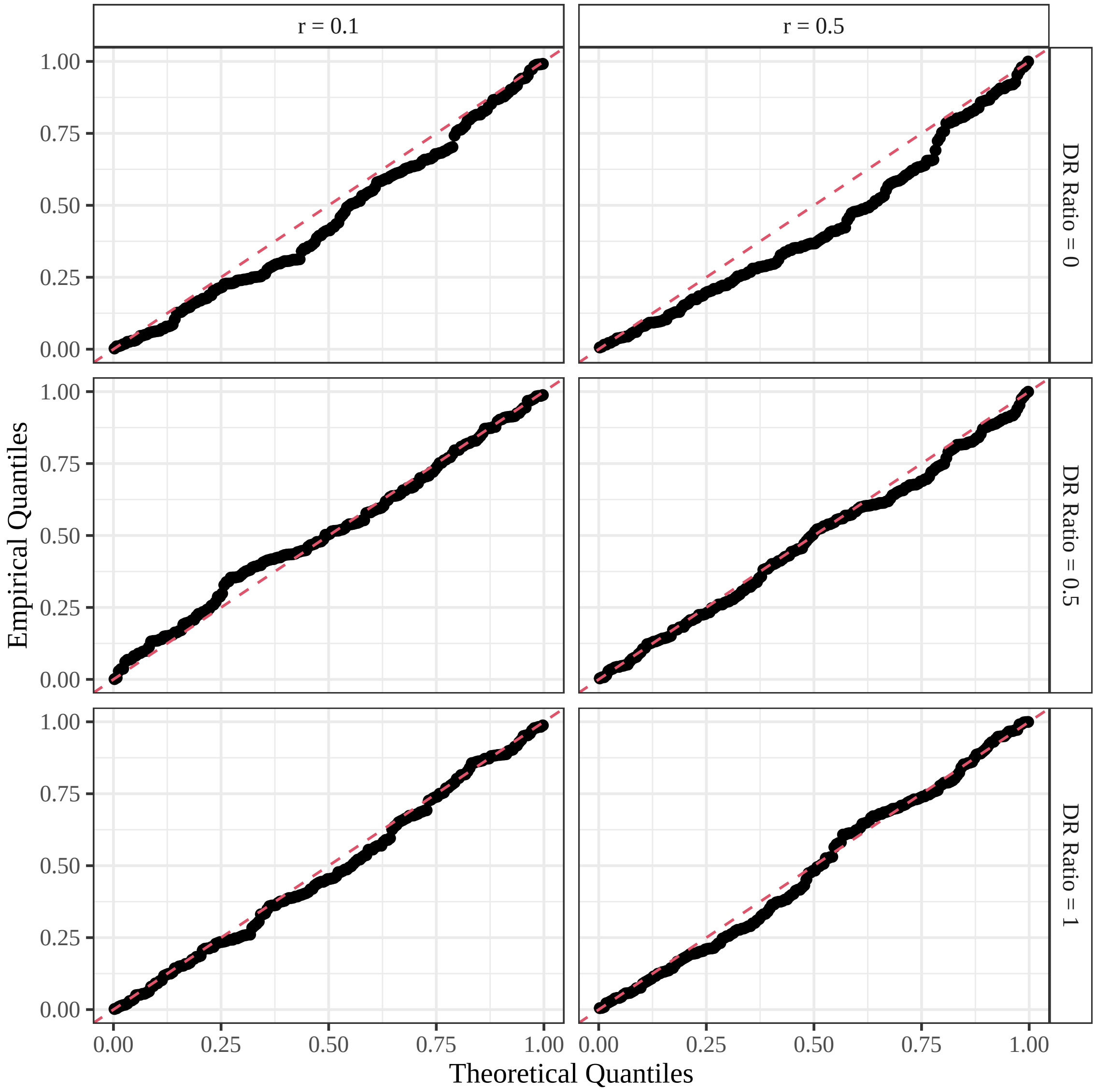


Figure S11: Quantile-quantile plots of the bootstrap approach's $p$-values (Section 2.5), testing against the null of equilibrium, when the sample size is $n = 25$ and the ploidy is 8, stratified by allele frequency (column facets) and ratio of the double reduction parameter to the maximum possible double reduction rate from Section S7.1 (row facets). These $p$-values were calculated in settings where the null was satisfied, and so should lie on the $y = x$ line (red-dashed line).

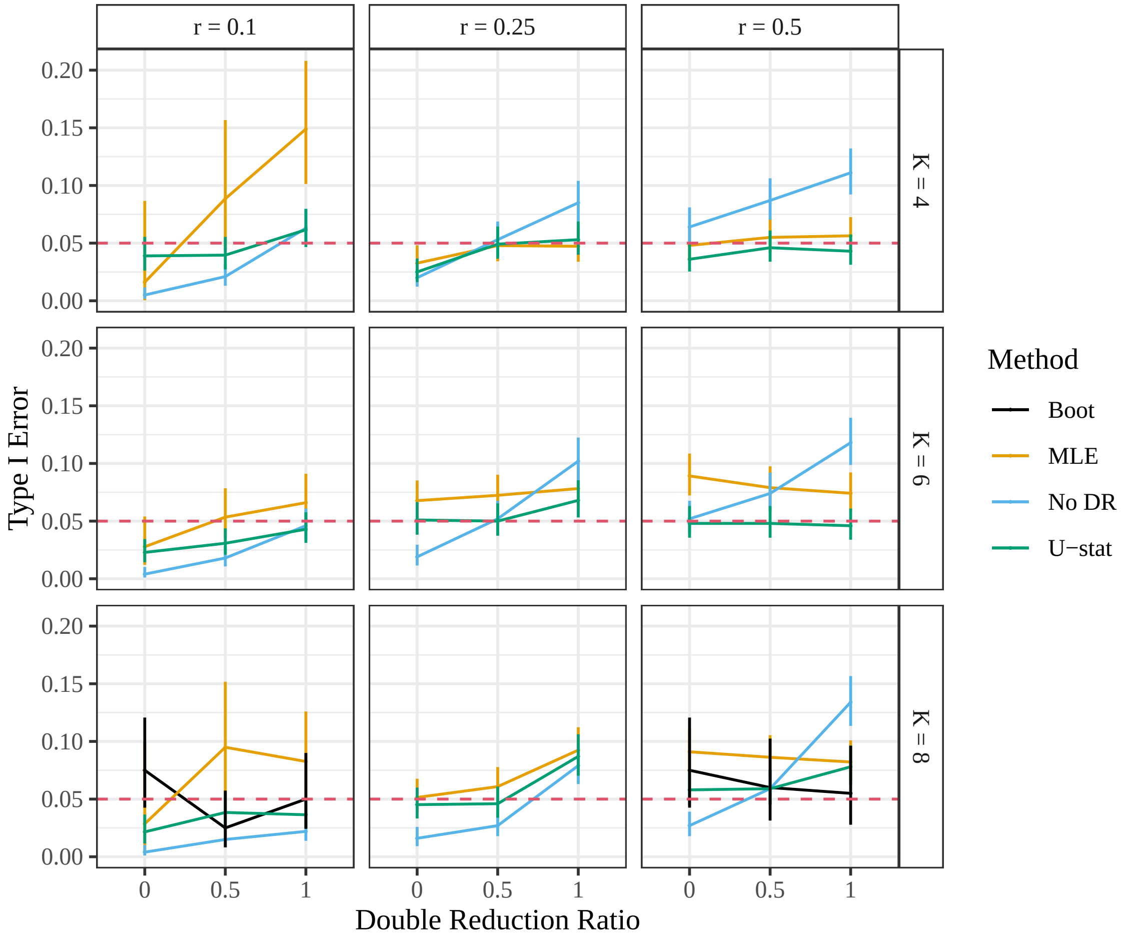


Figure S12: Type I error ($y$-axis) with corresponding 95% confidence intervals when testing against the null of equilibrium. Since equilibrium is satisfied and each test was run at a significance level of 0.05, the type I error should be near 0.05 (the red dashed line). Tests were run with a sample of size $n = 25$. Results are stratified by the ratio of double reduction to the maximum rate possible ($x$-axis), ploidy (row-facets), allele frequency (column-facets), and method (color). The $U$-statistic approach (Section 2.4) is in green, the likelihood approach (Section 2.3) is in orange, the bootstrap approach (Section 2.5) is in black, and the classical approach assuming no double reduction (Section S10) is in blue. The $U$-statistic, likelihood, and bootstrap approaches are at most slightly anti-conservative, while the classical approach only controls type I error when the double reduction ratio is 0.

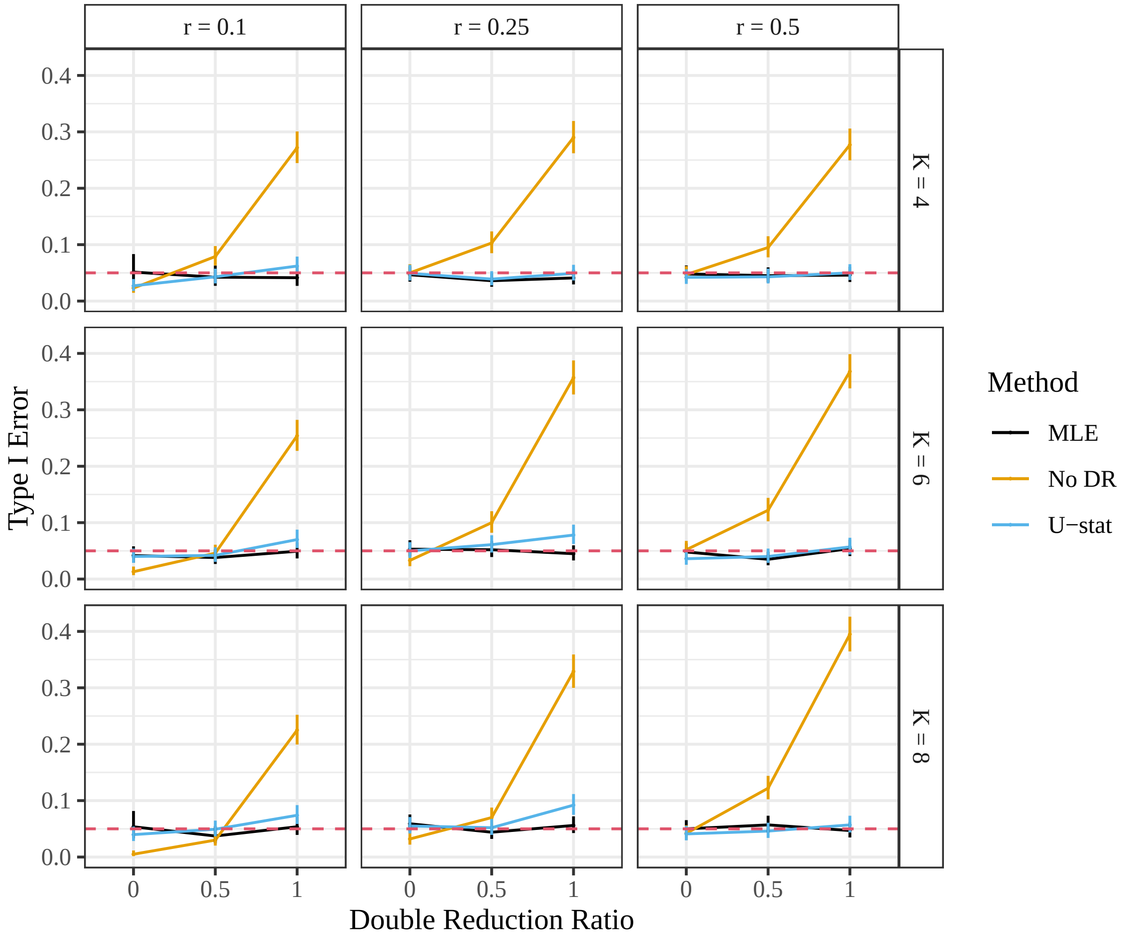


Figure S13: Type I error ($y$-axis) with corresponding 95% confidence intervals when testing against the null of equilibrium. Since equilibrium is satisfied and each test was run at a significance level of 0.05, the type I error should be near 0.05 (the red dashed line). Tests were run with a sample of size $n = 100$. Results are stratified by the ratio of double reduction to the maximum rate possible ($x$-axis), ploidy (row-facets), allele frequency (column-facets), and method (color). The $U$-statistic approach (Section 2.4) is in blue, the likelihood approach (Section 2.3) is in black, and the classical approach assuming no double reduction (Section S10) is in orange. The $U$-statistic and likelihood approaches are at most slightly anti-conservative, while the classical approach only controls type I error when the double reduction ratio is 0.

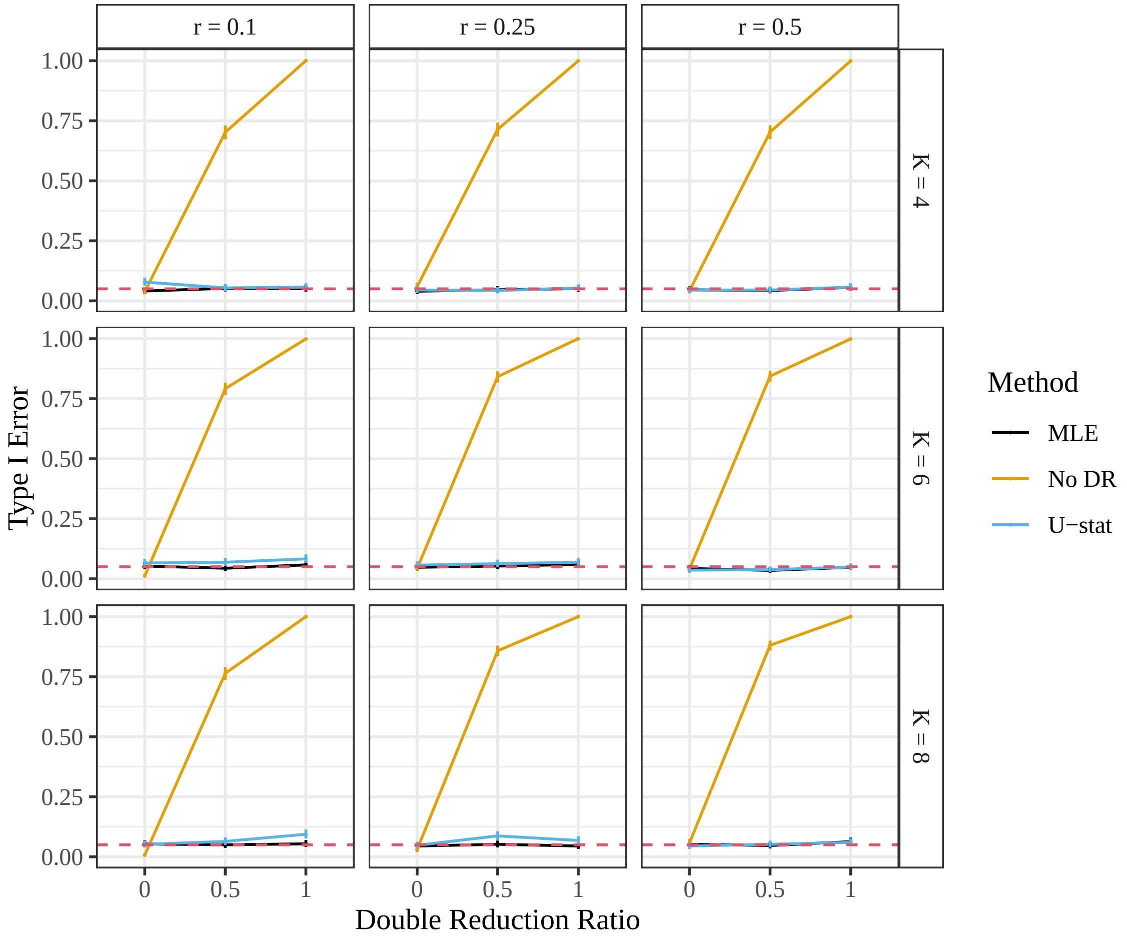


Figure S14: Type I error ($y$-axis) with corresponding 95% confidence intervals when testing against the null of equilibrium. Since equilibrium is satisfied and each test was run at a significance level of 0.05, the type I error should be near 0.05 (the red dashed line). Tests were run with a sample of size $n = 1000$. Results are stratified by the ratio of double reduction to the maximum rate possible ($x$-axis), ploidy (row-facets), allele frequency (column-facets), and method (color). The $U$-statistic approach (Section 2.4) is in blue, the likelihood approach (Section 2.3) is in black, and the classical approach assuming no double reduction (Section S10) is in orange. The $U$-statistic and likelihood approaches are at most slightly anti-conservative, while the classical approach only controls type I error when the double reduction ratio is 0.

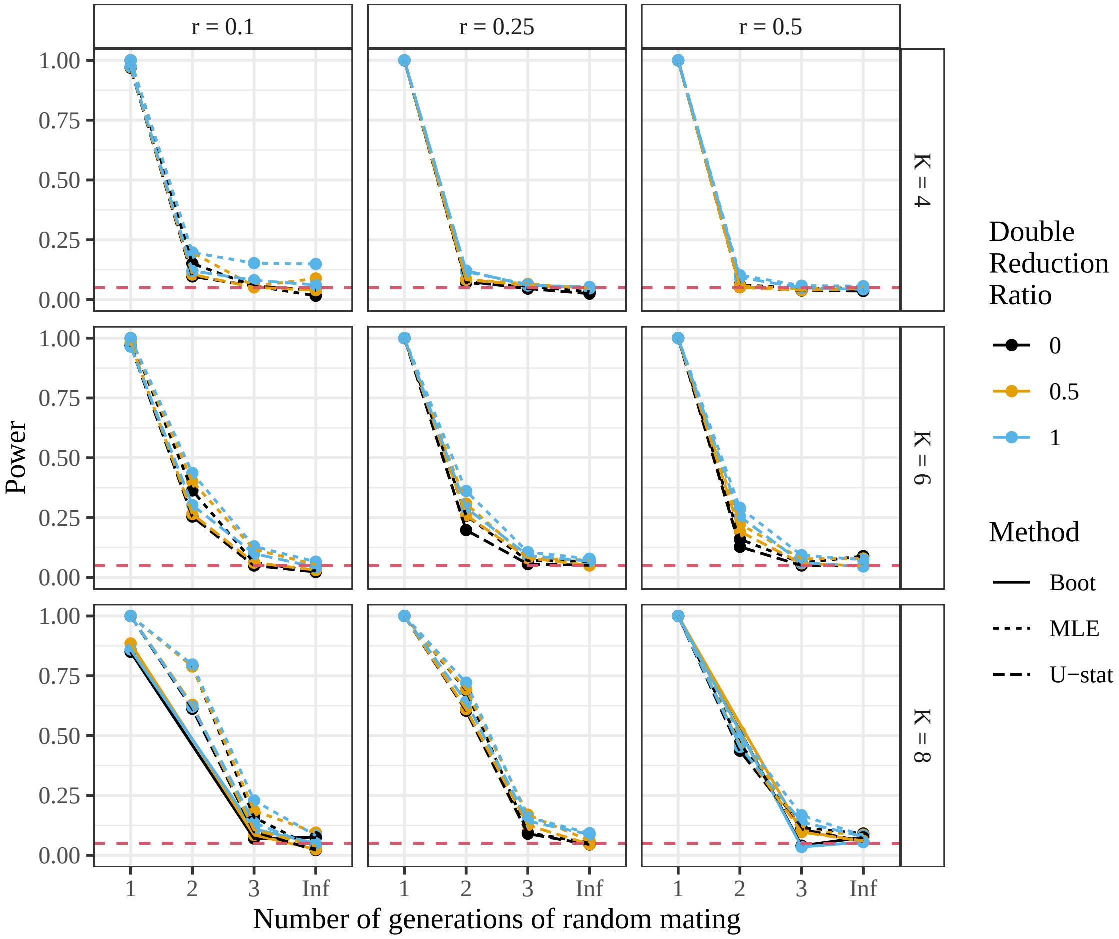


Figure S15: Power ($y$-axis) of the $U$-statistic approach (Section 2.4), the likelihood approach (Section 2.3), and the bootstrap approach (Section 2.5) when testing against the null of equilibrium. Results are stratified by the number of generations of random mating ($x$-axis), the ploidy (row facets), the allele frequency (column facets), and the ratio of double reduction to the maximum rate possible (color). The null is satisfied at “Inf” and, since each test is run at a 0.05 significance level, the “power” should be 0.05 in those scenarios. These are results for a sample of size $n = 25$.

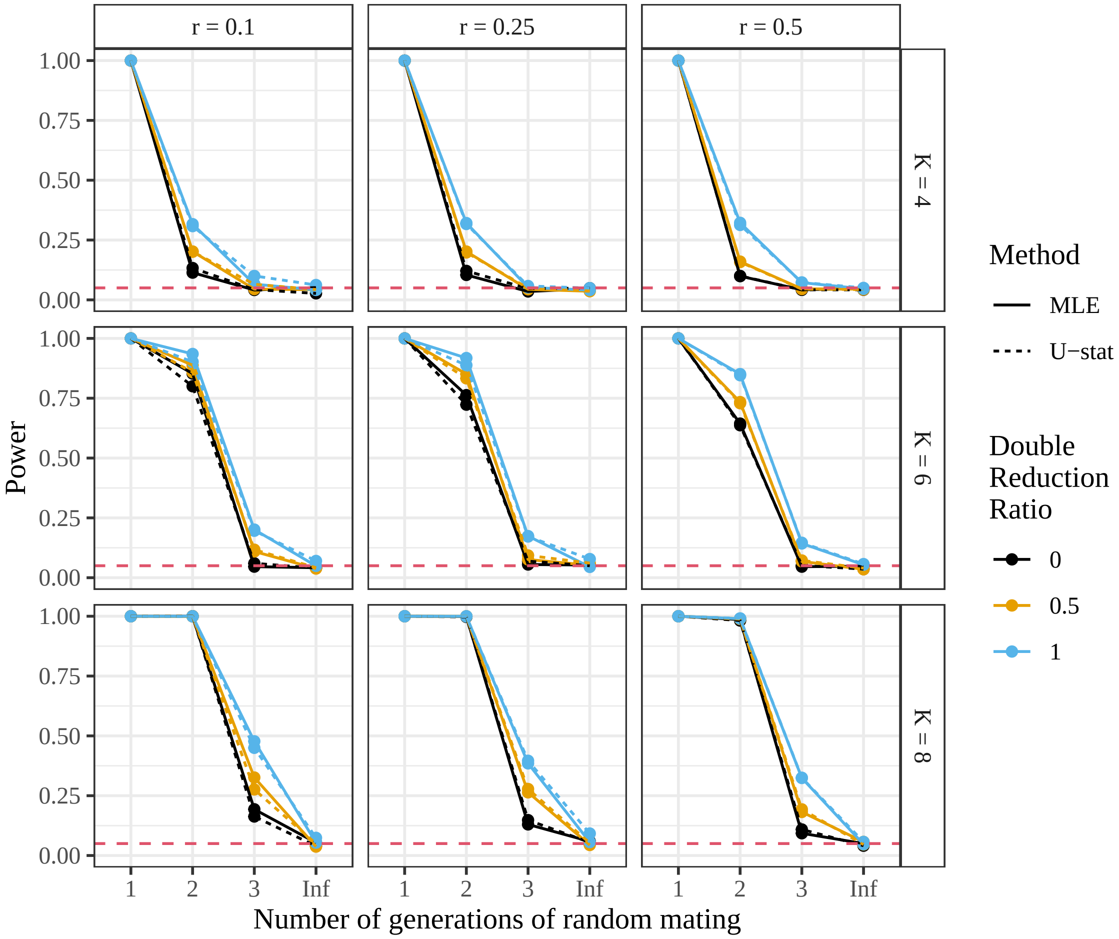


Figure S16: Power ($y$-axis) of the $U$-statistic approach (Section 2.4) and the likelihood approach (Section 2.3) when testing against the null of equilibrium. Results are stratified by the number of generations of random mating ($x$-axis), the ploidy (row facets), the allele frequency (column facets), and the ratio of double reduction to the maximum rate possible (color). The null is satisfied at "Inf" and, since each test is run at a 0.05 significance level, the "power" should be 0.05 in those scenarios. These are results for a sample of size $n = 100$.

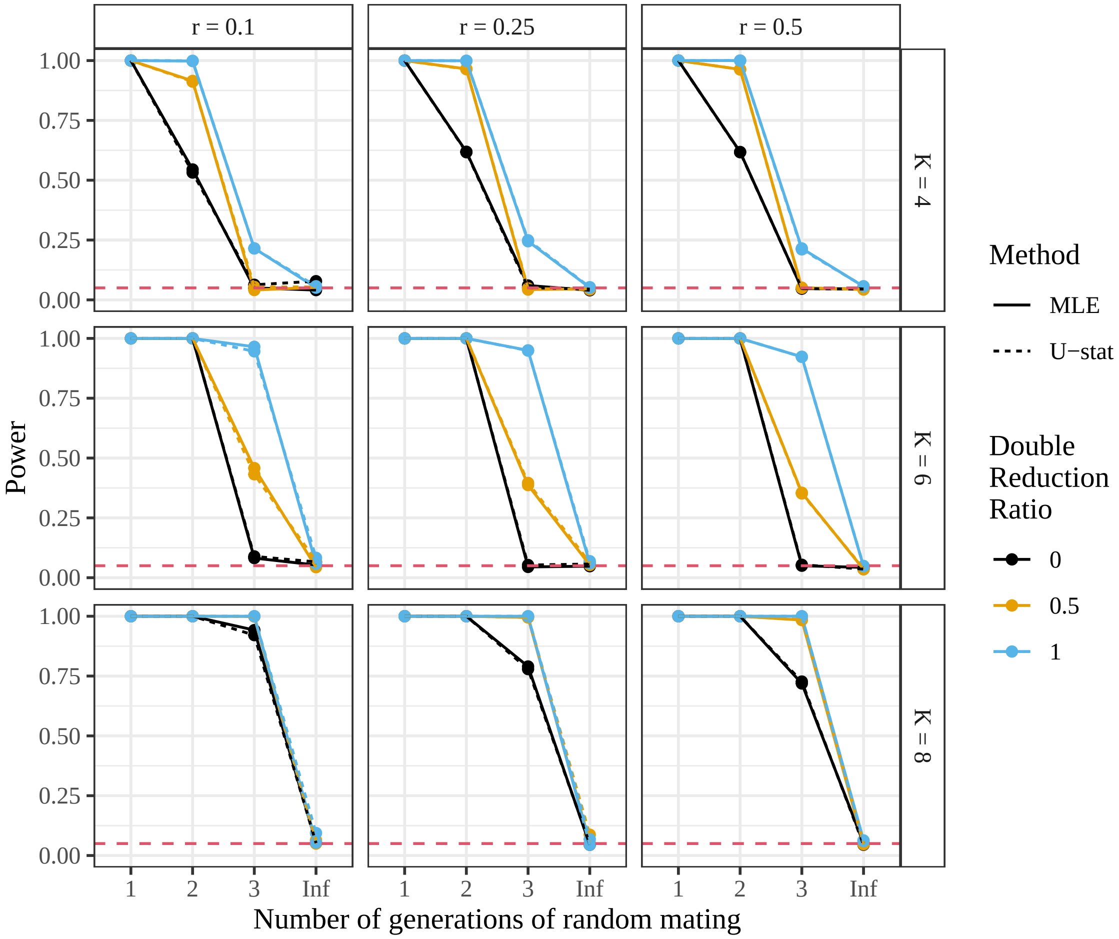


Figure S17: Power ($y$-axis) of the $U$-statistic approach (Section 2.4) and the likelihood approach (Section 2.3) when testing against the null of equilibrium. Results are stratified by the number of generations of random mating ($x$-axis), the ploidy (row facets), the allele frequency (column facets), and the ratio of double reduction to the maximum rate possible (color). The null is satisfied at "Inf" and, since each test is run at a 0.05 significance level, the "power" should be 0.05 in those scenarios. These are results for a sample of size $n = 1000$.

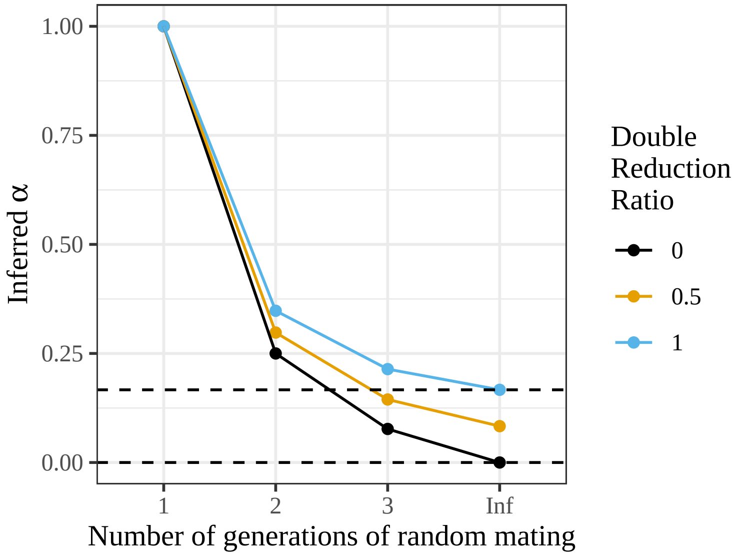


Figure S18: Inferred double reduction rate ($y$-axis) given gamete frequencies at each generation of random mating ($x$-axis) and the true double reduction rate (color). Results are similar for different values of the allele frequency $r$, and so only one scenario is shown. Only points within the dashed black lines are consistent with theoretical values of double reduction. Equilibrium is reached at "Inf".

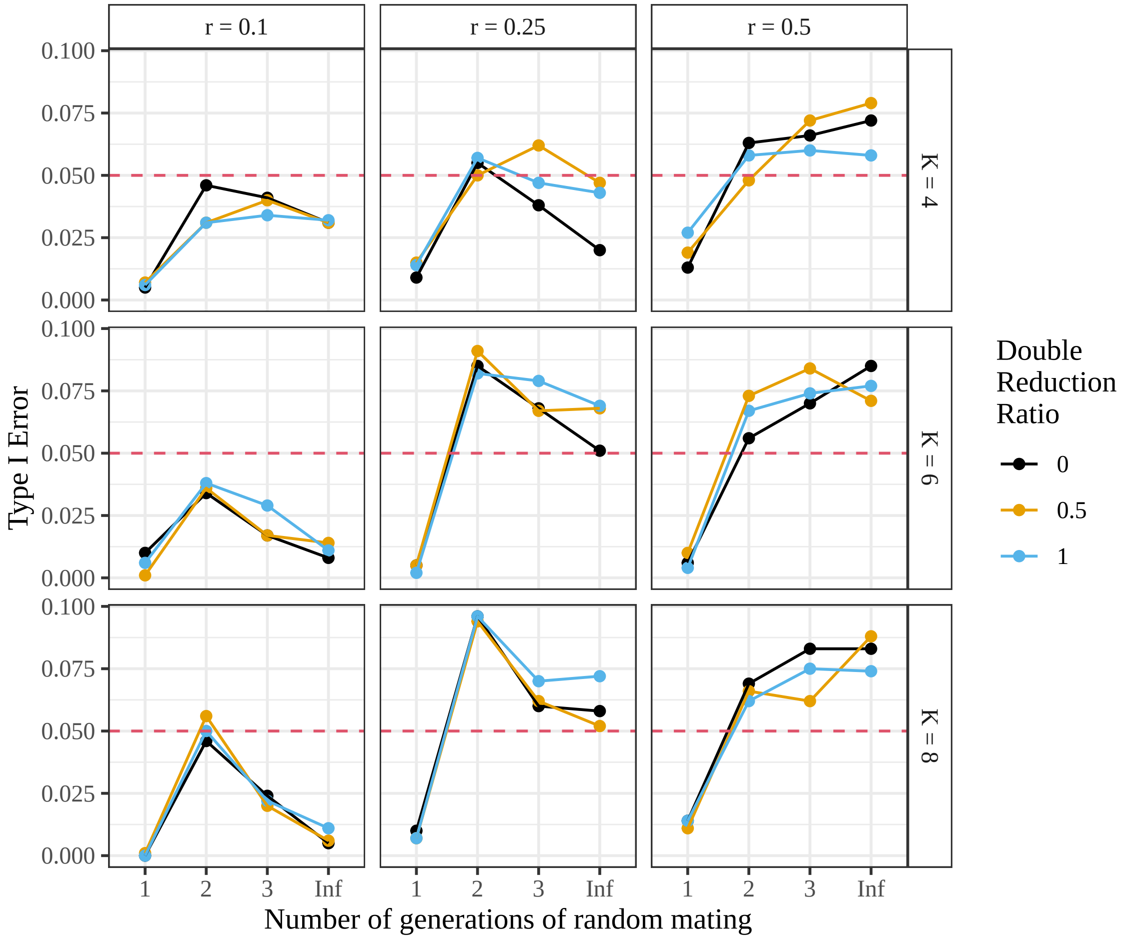


Figure S19: Type I error ($y$-axis) for the test of random mating, stratified by number of generations of random mating ($x$-axis), ploidy (row facets), allele frequency (column facets), and double reduction ratio (color), for a sample of size $n = 25$. All scenarios are null, and the test is sometimes slightly anti-conservative.

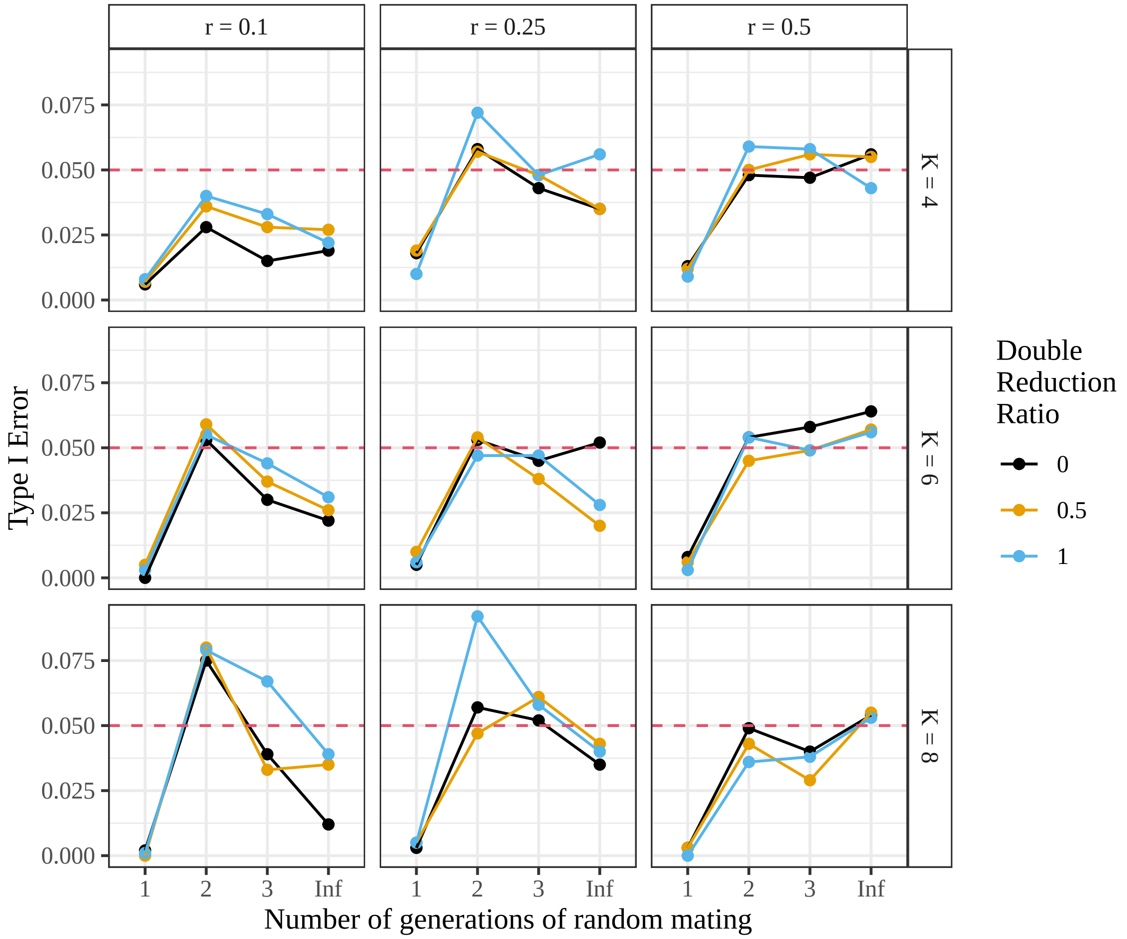


Figure S20: Type I error ($y$-axis) for the test of random mating, stratified by number of generations of random mating ($x$-axis), ploidy (row facets), allele frequency (column facets), and double reduction ratio (color), for a sample of size $n = 100$. All scenarios are null, and the test is mostly able to control Type I error.

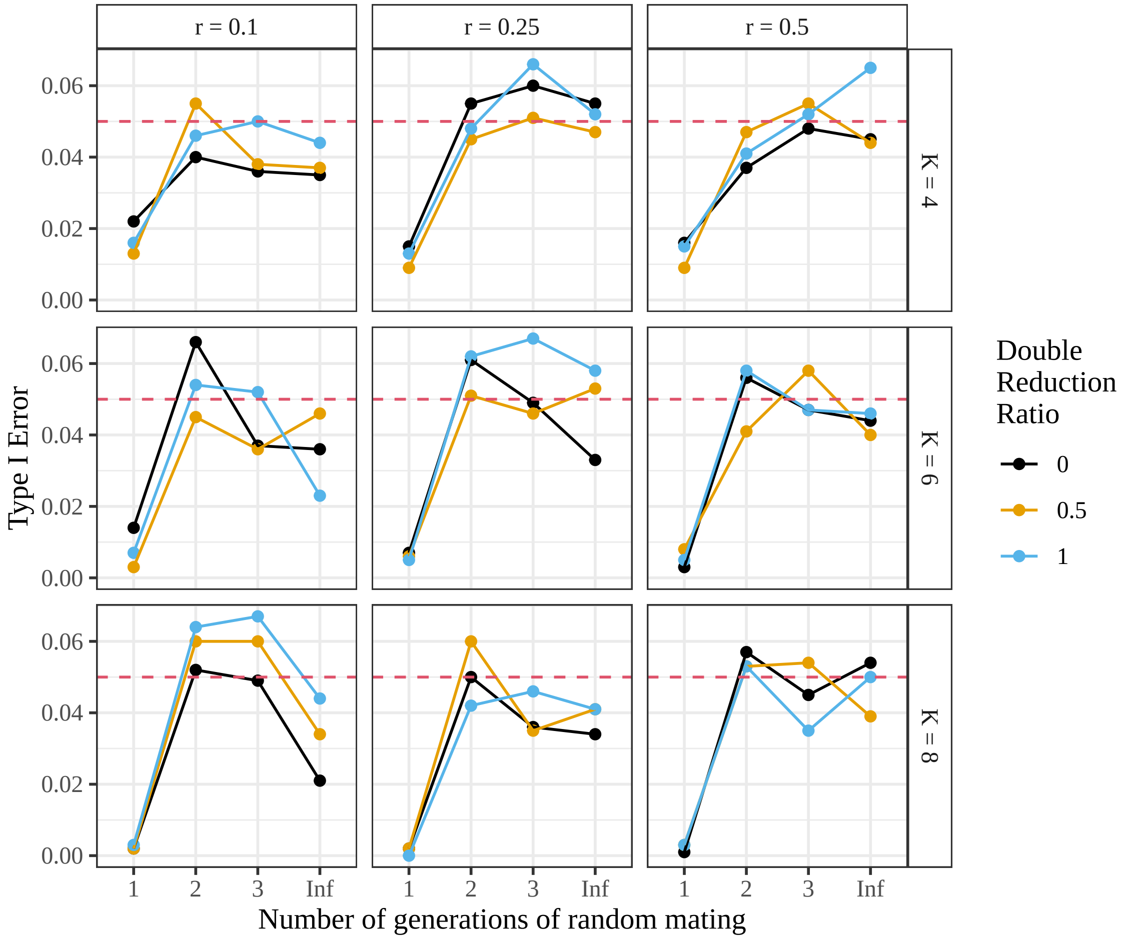


Figure S21: Type I error ($y$-axis) for the test of random mating, stratified by number of generations of random mating ($x$-axis), ploidy (row facets), allele frequency (column facets), and double reduction ratio (color), for a sample of size $n = 1000$. All scenarios are null, and the test is mostly able to control Type I error.

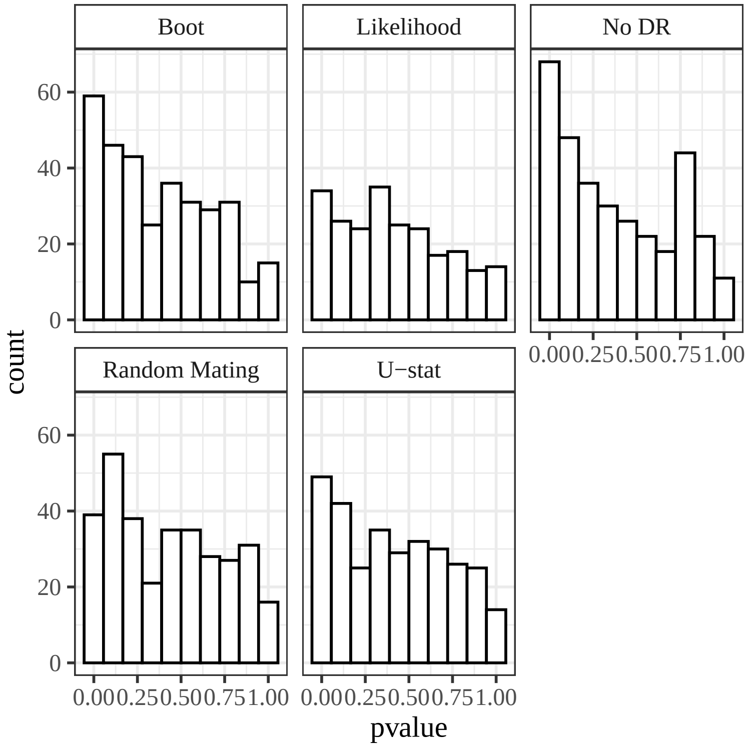


Figure S22: Histogram of $p$-values from the Sturgeon data of Delomas et al. [2021]. The facets stratify the different tests: the likelihood ratio test for equilibrium (Likelihood) (Section 2.3), the $U$-statistic test for equilibrium (U-stat) (Section 2.4), the likelihood ratio test for equilibrium that assumes no double reduction (No DR) (Section S10), the bootstrap procedure for equilibrium (Boot) (Section 2.5), and the likelihood ratio test for random mating (Random Mating) (Section 2.2).

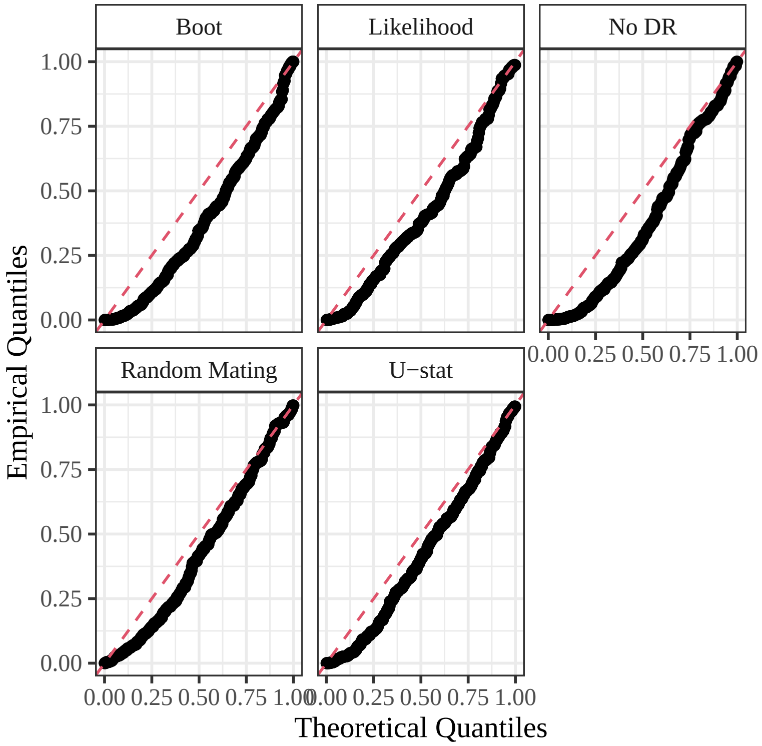


Figure S23: QQ-plots against the uniform distribution of the $p$-values from the Sturgeon data of Delomas et al. [2021]. The facets stratify the different tests: the likelihood ratio test for equilibrium (Likelihood) (Section 2.3), the $U$-statistic test for equilibrium (U-stat) (Section 2.4), the likelihood ratio test for equilibrium that assumes no double reduction (No DR) (Section S10), the bootstrap procedure for equilibrium (Boot) (Section 2.5), and the likelihood ratio test for random mating (Random Mating) (Section 2.2).

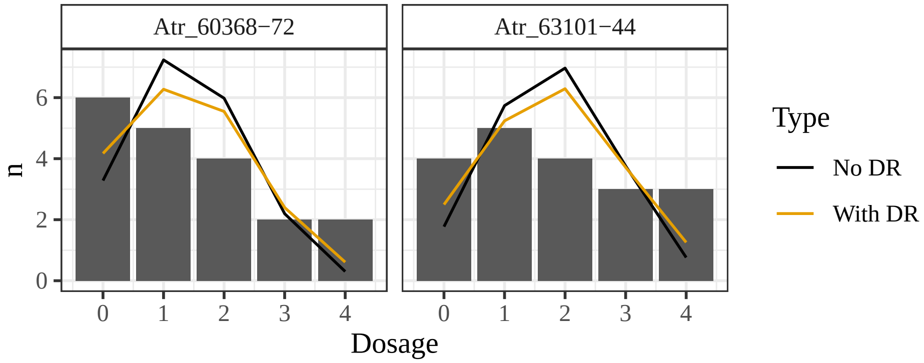


Figure S24: Genotype counts (bars) and expected counts (lines) when assuming either no double reduction or double reduction (color) for two loci (facets) from the Sturgeon data that have small $p$-values when one assumes no double reduction and large $p$-values when one allows for double reduction.

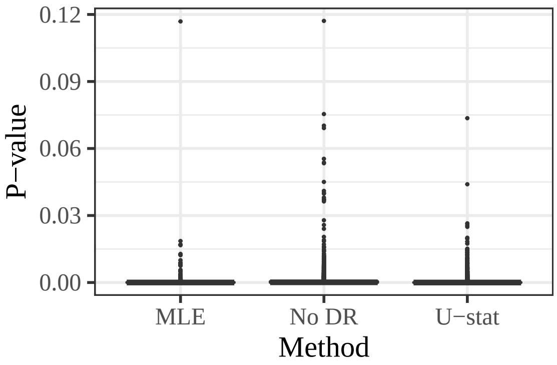


Figure S25: Boxplots of $p$-values ($y$-axis) for each method ($x$-axis) fit on the data from Shirasawa et al. [2017] (Section 3.3). Methods considered are the likelihood approach (MLE) (Section 2.3), the $U$-statistic approach (Section 2.4) ($U$-stat), and the naive approach (Section S10) (No DR). Data are not at equilibrium, so all $p$-values should be near 0.

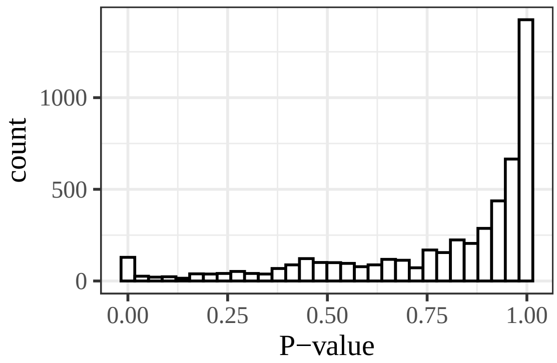


Figure S26: Histogram of $p$-values from the test for random mating (Section 2.2) applied to the data from Shirasawa et al. [2017] (Section 3.3). Random mating is fulfilled, so ideally we should see a uniform distribution of $p$-values. The test is conservative, likely because of the large number of genotypes that have 0 counts.

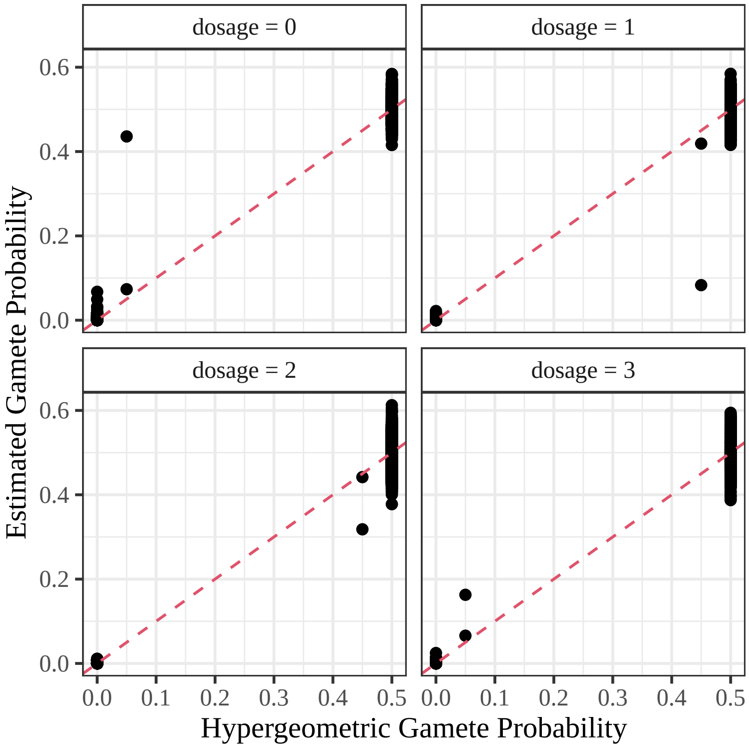


Figure S27: Estimated gamete probabilities using the random mating model of Section 2.2 ($y$-axis) plotted against the theoretical hypergeometric probabilities given the estimated parental genotype ($x$-axis). The dashed line is the $y = x$ line.

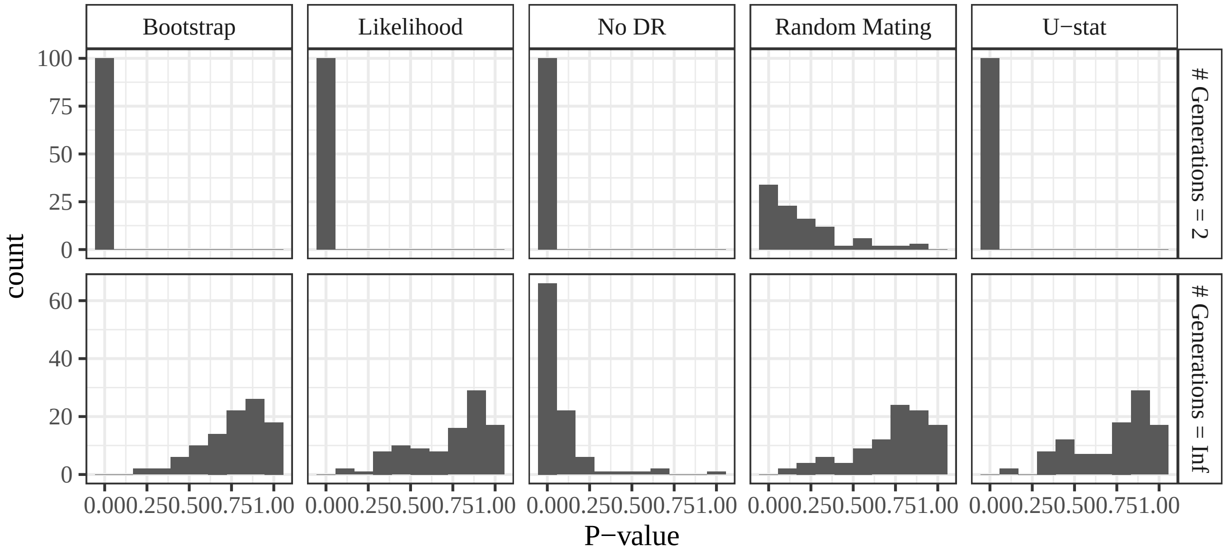


Figure S28: Histograms of $p$-values for each method (column facets) after two or infinitely many generations of random mating (row facets). Equilibrium is fulfilled under infinitely many generations of random mating, but is not fulfilled after only two generations. The random mating hypothesis is fulfilled in both scenarios. The methods explored, in order from left to right, are the bootstrap procedure (Section 2.5), the likelihood approach (Section 2.3), the approach that does not account for double reduction (Section S10), the test for random mating (Section 2.2), and the $U$-statistic approach (Section 2.4)
